# Spatial profiling of neuro-immune interactions in gastroenteropancreatic NETs

**DOI:** 10.1101/2023.07.01.547336

**Authors:** Suzann Duan, Travis W. Sawyer, Brandon L. Witten, Heyu Song, Tobias Else, Juanita L. Merchant

## Abstract

Gastroenteropancreatic neuroendocrine tumors (GEP-NETs) are heterogeneous malignancies that arise from complex cellular interactions within the tissue microenvironment. Here, we sought to decipher tumor-derived signals from the surrounding microenvironment by applying Nanostring Digital Spatial Profiling (DSP) to hormone-secreting and non-functional GEP-NETs. DSP was used to evaluate the expression of 40 neural and immune-related proteins in surgically resected duodenal and pancreatic NETs (n=20) primarily comprised of gastrinomas (18/20). A total of 279 regions of interest were examined between tumors, adjacent normal and abnormal-appearing epithelium, and the surrounding stroma. The results were stratified by tissue type and *Multiple Endocrine Neoplasia I (MEN1)* status and protein expression was validated by immunohistochemical (IHC) staining. A tumor immune cell autonomous inflammatory signature was further evaluated by IHC and RNAscope, while functional pro-inflammatory signaling was confirmed using patient-derived duodenal organoids. Gastrin-secreting and non-functional pancreatic NETs showed a higher abundance of immune cell markers and immune infiltrate compared to duodenal gastrinomas. Tumors displayed strong intra-tissue variation in the expression of neural- and immune-related proteins. Compared to non-*MEN1* tumors, *MEN1* gastrinomas showed reduced expression of immune cell markers and upregulated expression of neuropathological proteins. Duodenal gastrinomas showed strong expression of the pro-inflammatory and pro-neural factor IL-17B. Treatment of human duodenal organoids with IL- 17B activated NF-kB and STAT3 signaling and induced the expression of neuroendocrine markers. In conclusion, multiplexed spatial protein analysis identified tissue-specific neuro- immune signatures in GEP-NETs. Moreover, duodenal gastrinomas cell autonomously express immune and pro-inflammatory factors, including tumor-derived IL-17B, that stimulate the neuroendocrine phenotype.

## Introduction

Despite rising incidence and prevalence rates, the mechanisms that underlie gastroenteropancreatic tumors (GEP-NETs) remain poorly understood. GEP-NETs comprise a complex group of cancers that display diverse molecular features, mixed differentiation status, and broad proliferative potential (Kawasaki *et al*. 2018). GEP-NETs that secrete bioactive hormones are classified as functional, whereas non-hormone secreting tumors are considered non-functional. Gastrinomas are functional GEP-NETs that secrete gastrin peptide and most commonly arise in the pancreas, proximal duodenum, and gastric antrum comprising the “gastrinoma triangle” (Stabile *et al*. 1984). Among these, duodenal gastrinomas (DGASTs) associated with the *Multiple Endocrine Neoplasia I (MEN1)* Syndrome are the most malignant, presenting as small (70% <1.5cm), multicentric (90%) tumors that are more likely to metastasize, with 30% exhibiting lymph node metastases upon diagnosis (Jensen & Norton *et al*. 2017). Despite their association with clinically aggressive metrics, the signals that direct the emergence of *MEN1* gastrinomas remain enigmatic.

Prior efforts to stratify GEP-NETs for treatment have largely centered on bulk sequencing of tumor biopsies and surgically resected tissues to identify genetic and epigenetic alterations (Jiao *et al*. 2011, How-Kit *et al*. 2015, Scarpa *et al*. 2017, Panarelli *et al*. 2019, Pucini et al. 2020, Melone *et al*. 2022, Wu *et al*. 2022). Our previous transcriptomic analysis of a small cohort of human DGASTs identified expression of pro-inflammatory cytokines in the tumor and preneoplastic Brunner’s glands, indicating a role in neoplastic transformation (Rico *et al*. 2021). In these studies, DGASTs were shown to express functional pro-inflammatory cytokines, including TNF-11, that promoted the neuroendocrine phenotype in organoid culture studies. Importantly, activation of inflammatory signaling was immune cell-independent, as expression of these cytokines and downstream signaling was restricted to endocrine cells and the tumor stroma rather than the tumor immune compartment (Rico *et al*. 2021). Consistent with these observations, prior immuno-typing of GEP-NETs showed that a majority of grade 1 and 2 (G1/2) tumors exhibit an immune excluded or immunosuppressive microenvironment compared to G3 NETs and neuroendocrine carcinomas (NECs) (Sampedro-Nunez *et al*. 2018, Ferrata *et al*. 2019, Bosch *et al*. 2019, Puccini *et al*. 2020, Busse *et al*. 2020). These studies and others suggest that GEP-NETs may be clinically stratified by their microenvironments, including the enrichment of immune infiltrates and cancer-associated fibroblasts (Lou *et al*. 2022). Thus, further subtyping of GEP-NETs by immune and stromal status may elucidate approaches to therapeutically target the tumor microenvironment and unlock the potential for personalized medicine.

Until recently, the absence of reliable methods to unmix tumor- and stroma-derived signals has precluded a comprehensive understanding of how GEP-NETs arise in different tissues. Advances in spatial profiling technologies have addressed these limitations by illuminating cellular interactions in the tumor microenvironment with astonishing detail (Brady *et al*. 2021, Yang *et al*. 2022, Nino *et al*. 2022, Carter *et al*. 2023, Kim *et al*. 2023). Here, we applied digital spatial profiling to 18 gastrin-secreting and 2 non-functional GEP-NETs and elucidated their unique molecular features that provided an indication of their cellular origins. By combining this approach with *in vitro* studies in human-derived organoids, we demonstrate the convergence of cell autonomous immune and pro-inflammatory proteins that suggests their role in neuroendocrine differentiation and tumorigenesis.

### Study Objective

Perform Nanostring digital spatial profiling on gastrin-secreting and non-functional GEP-NETs to identify neuro-immune profiles in the tumor microenvironment.

## Materials and Methods

### Human Tissue Collection

De-identified formalin-fixed paraffin-embedded (FFPE) sections of surgically resected GEP-NETs were collected under IRB approval #HUM00115310 from the Department of Pathology at the University of Michigan and the Department of Endocrinology at the University of Pennsylvania. Fresh normal-appearing duodenal mucosa was collected during Whipple procedures through the Tissue Acquisition and Mouse Resource Core (TACMASR) at the University of Arizona Comprehensive Cancer Center.

### Nanostring GeoMx Digital Spatial Profiling

Five-micron FFPE tissue sections were baked at 65L for 60 min prior to deparaffinization in xylene and rehydration in 100%, 90%, and 70% ethanol. Slides were washed in Tris Buffered Saline (TBS), then subjected to heat-mediated antigen retrieval using sodium citrate buffer (pH 6.0). Slides were incubated in a pressure cooker for 15 min and allowed to cool to 24L before proceeding with blocking and immunostaining steps. To identify regions of interest (ROIs) for protein expression analysis, tumor sections were stained using the Nanostring Tumor Microenvironment Morphology markers: PanCK (pan cytokeratin denoting epithelial cells, CD45 (immune cells) and SYTO13 (a DNA-intercalating dye marking cell nuclei).

Expression analysis of 40 neural- and immune-related proteins was performed using the Nanostring Neural Cell Profiling Core supplemented with the Alzheimer’s Disease Module and Parkinson’s Disease Module (Nanostring Technologies, Seattle, WA). The Neural Cell Profiling Core and Modules are comprised of a multiplex panel of antibodies conjugated to ultraviolet light (UV)-cleavable oligomers that enable the detection and quantitation of spatially resolved protein expression. The Nanostring GeoMx Digital Spatial Profiler instrument was used to acquire a high resolution wide-field scan of each tissue section from which ROIs (200 micron by 200 micron) were selected for UV-activated oligo collection. Collected oligos were hybridized to unique probes and pooled for analysis using the Nanostring nCounter instrument, as described by the manufacturer.

The first DSP study was performed on a discovery cohort consisting of 4 surgically resected duodenal gastrinomas (DGAST), 3 pancreatic gastrinomas (PGAST), 2 non-functional pancreatic NETs (NF-PNET). Duodenal mucosa and pancreas from non-tumor bearing subjects were also included as a comparison. Of these, the four duodenal gastrinoma-bearing patients were initially described in a previous study (Rico *et al*. 2021). Eight ROIs were selected per slide for a total of 96 ROIs. The second DSP study was performed on an expanded cohort of 12 different surgically resected duodenal gastrinomas. Of the 216 ROIs selected, 183 passed quality control parameters and were included for analysis. These included 47 normal BGs, 76 abnormal BGs, and 60 Tumor ROIs across the 12 patients. A series of quality control, normalization, and background subtraction steps were performed according to published Nanostring recommendations. Background subtraction was performed using the average of two IgG Isotype controls and protein expression values were normalized to the mean of three housekeeping proteins included in the antibody panel: GAPDH, Histone H3, and S6. Linear Mixed Modeling and Benjamini-Hochberg multiple test correction was applied to test for statistically significant differences.

### DSP Data Visualization

Data analysis and visualization including heatmaps and PCA were completed with Python 3 using the “seaborn” and “pandas” analysis packages. Prior to visualization, spreadsheets containing raw and normalized expression reads for each extracted ROI were loaded into the program as a dataframe. Markers used for normalization including GAPDH, Histone H3, S6, Mouse IgG1, Mouse IgG2a, and Rabbit IgG were removed from the dataframe, and a unique tissue class identifier was assigned to each ROI. Each dataframe was transformed to express the z-score of the individual proteins relative to the entire sample set. Heatmaps were generated by plotting the z-scores for each marker for each of the tissue groups using the seaborn’s built-in heatmap function.

Principal component analysis (PCA) was conducted on each dataset after z-score transformation, with no dimension reduction applied. PCA plots were generated by plotting the first and second principal components for each ROI on two orthogonal axes, and color coding according to the tissue type. PCA biplots were generated using a custom function to plot the projection of specific markers into PCA space by extracting the first and second loading coefficients. These coefficients were used to plot lines superimposed on the PCA plots, which point in the direction of variance corresponding to each marker.

Unsupervised clustering was conducted on the datasets using the seaborn built-in clustermap function applied to each normalized data frame. The data was clustered using unsupervised hierarchical clustering for both axes, using the Euclidean distance metric (2-norm), and linkages calculated using the UPGMA algorithm (Sokal & Michener, 1958). Dendrograms were then superimposed on the heatmaps to visualize clustering.

### Immunohistochemical and Immunofluorescent Staining

Five-micron FFPE sections were baked at 65L for 60 min and then deparaffinized in xylene three times for 5 min. Tissues were rehydrated in a series of 100%, 90%, and 70% ethanol washes, and then rinsed with Tris Buffered Saline (TBS) for 5 min. Heat-mediated antigen retrieval was performed by heating slides in Tris-EDTA buffer (pH 9.0) (Sakura Finetek, Torrance, CA) for 30 min at 95L. Slides were allowed to cool to 24L for 15 min, and then rinsed for 1 min under running distilled water before washing three times in TBS. Sections were blocked with 5% bovine serum albumin (BSA) in TBS with 0.05% Tween-20 (TBST) for 1h at 24L. Next, slides were incubated in primary antibodies overnight (16h) at 4L. Slides were washed three times in TBST, and then incubated in anti-Mouse IgG or anti-Rabbit IgG secondary antibodies conjugated to Horse Radish Peroxidase (ImmPACT DAB, Vector Laboratories, Newark, CA) in a humid chamber for 30 min at 24L. Slides were washed, and then incubated in DAB substrate (ImmPACT DAB kit, Vector Laboratories) for 30-60 sec or until the chromogenic stain could be visualized. Slides were immediately washed in running distilled water for 1 min prior to counterstaining cell nuclei with diluted Gill’s Hematoxylin (Leica Biosystems, Wetzlar, Germany). Slides were washed under running distilled water, briefly dipped in Bluing Reagent (Leica Biosystems) prior to washing again and dehydrating in a series of 70%, 90%, and 100% ethanol. Slides were washed in xylene three times for 5 min and then mounted with Paramount mounting medium.

For immunofluorescent staining of FFPE slides, sections were processed as described previously and blocked in 10% donkey serum in 1% BSA TBST for 1 h at 24. Sections were then incubated in the following primary antibodies overnight at 4L: IL-17B (1:100, Cat #1248, RRID:AB_354696, R&D Systems, Minneapolis, MN), TNF-alpha (1:100 Cat #AF-410, RRID:AB_354479, R&D Systems), and phosho-STAT3 Tyr705 (1:100, Cat #9145, RRID:AB_2491009, Cell Signaling Technologies). Slides were washed three times in TBST and then incubated in Alexa Fluor-conjugated secondary antibodies diluted 1:500 in TBST with 1% BSA (Thermo Fisher Scientific, Waltham, MA). Slides were washed three times in TBS for 5 min at 24L prior to mounting with coverslips and Prolong Gold anti-fade mounting medium with 4’, 6-diamidino-2-phenylindole (DAPI) (Thermo Fisher Scientific). IHC and IF stained slides were imaged using the Olympus BX53F epifluorescence microscope (Center Valley, PA).

For immunostaining of duodenal organoids, the treated organoids were washed twice with pre-warmed Dulbecco’s PBS (DPBS). Organoids were fixed for 20 min at 24L in 4% PFA that was pre-warmed to 37L. Organoids were permeabilized in 0.5% Triton X-100 for 15 min at 24L. Organoids were washed twice with DPBS and then blocked in 10% donkey serum and 1% BSA for 30 min at 24L. Organoids were incubated in the following primary antibodies for 3 h at 24L: gastrin (1:1000, Cat #A0568, A0568, AB_2757270, Dako Agilent Technologies), Chromogranin A (1:100, Cat #ab45179, Abcam, RRID:AB_726879, Cambridge, UK), and Synaptophysin (1:400, Cat #9020, RRID:AB_2631095, Cell Signaling Technologies).

### Organoid Culture

Human duodenal organoid lines were generated from normal-appearing duodenal biopsies from two patients undergoing Whipple procedure (Rico *et al*. 2021). Organoids lines were subcultured in complete organoid media as described previously (Supplementary Table 1). Media was supplemented with 10 µM SB202190, a selection p38 MAP kinase inhibitor and 10 nM nicotinamide (Tocris Bioscience, Bristol, UK) for the first day of passaging to prevent anoikis. Organoids were used for studies between passage numbers 5 and 10. Within 5–7 days post-seeding, duodenal organoids were treated with 0, 10, 25, 50, and 100 ng/mL of recombinant human IL-17B (R&D Systems, Minneapolis, MN) for 48 h and processed for downstream RNA expression analysis. For whole cell protein analysis, organoids were treated with 50 ng/mL of IL- 17B for 24 h, and a time-course treatment was conducted over 0, 0.5, 1, 2, 3, and 4 h to examine nuclear and cytoplasmic protein shuttling.

### Western Blot

Following 4 h of IL-17B treatment, duodenal organoids were scraped from wells and collected in ice-cold PBS. Organoids were dissociated from the Matrigel (Corning, Corning, NY) by gently pipetting ten times and centrifuging for 5 min at 300 x g at 4L. The Matrigel-containing supernatant was carefully aspirated off and the organoid pellet was resuspended in 1 mL ice-cold PBS. After mixing ten times, the organoids were centrifuged for 5 min at 300 x g at 4 and the supernatant was discarded. To generate whole cell extracts, the pellet was resuspended in 200 µL of ice-cold RIPA buffer (Thermo Fisher Scientific) supplemented with 1X HALT protease and phosphatase inhibitor (Thermo Fisher Scientific). Organoids were lysed by passing through a 20G syringe ten times and vortexing. Following 40 min incubation on ice, organoid lysates were centrifuged for 15 min at 15,000 x g at 4L and the supernatant was collected as the protein extract.

For subcellular fractionation, organoid pellets were resuspended in 160 µL of ice-cold hypotonic cytoplasmic lysis buffer comprised of 20 mM HEPES (pH 8.0), 0.1% IGEPAL, 1 mM dithiothreitol (DTT) and 1X HALT protease and phosphatase inhibitor prepared in Dulbecco’s PBS. Organoids were mixed by pipetting 20X and incubating on ice for 10 to 15 min. After the incubation, organoids were pulse vortexed, and then centrifuged for 5 min at 15,000 x g at 4. The supernatant was collected as the cytoplasmic protein fraction. The pellet was washed three times with cytoplasmic lysis buffer, then resuspended in 40 uL of a hypertonic nuclear lysis buffer comprised of 20 mM HEPES (pH 8.0), 20% glycerol, 0.2 mM EDTA (pH 8.0), 500 mM NaCl, 1.5 mM MgCl_2_, 0.1% IGEPAL, 1mM DTT, and 1X HALT protease and phosphatase inhibitor prepared in Dulbecco’s PBS. The pellet was vigorously vortexed and incubated on ice for 40 min. The solution was centrifuged for 15 min at 15,000 x g at 4L to obtain the nuclear protein fraction.

Ten micrograms of protein was loaded onto a 4-12% Bis-Tris precast NuPage Gel (Thermo Fisher Scientific). Gel electrophoresis was performed at 140 V for 1 h. Proteins were transferred for 7 min at 22-25 V onto a PVDF membrane using the iBlot2 transfer system (Thermo Fisher Scientific). Membranes were blocked with 5% BSA TBST for 1h at 24 prior to incubating with STAT3 (Cat #12640, RRID:AB_2629499, Cell Signaling Technologies) or NF- kB (p65) (Cat #8242, RRID:AB_10859369, Cell Signaling Technologies) antibodies diluted 1:1000 in blocking buffer overnight at 4. Membranes were washed three times with TBST, and then incubated for 1 h at 24L in HRP-conjugated anti-Rabbit IgG secondary antibody (Cat #7074, RRID:AB_2099233, Cell Signaling Technologies) diluted 1:3,000 in blocking buffer (Cell Signaling Technologies, Danvers, MA). Membranes were washed three times with TBST and incubated in ECL substrate solution (Thermo Fisher Scientific) for 1 min prior to developing images on film. Membranes were reprobed using Cell Signaling antibodies for Histone H3 (1:10,000, Cat #12648, RRID:AB_10544537), beta-tubulin (1:2,000, Cat #5346, RRID: AB_1950376), or GAPDH (1:2,000, Cat #5147, RRID:AB_10622025) as respective nuclear, cytoplasmic, and whole cell lysate loading controls.

### Quantitative Polymerase Chain Reaction (RT-qPCR)

RNA was extracted from duodenal organoids using the ReliaPrep miRNA Cell and Tissue Miniprep System following the manufacturer’s instructions (Promega, Madison, Wisconsin). Up to 1 microgram of cDNA was synthesized using SuperScript VILO after treatment with DNAse I to remove genomic DNA. Quantitative real-time PCR (RT-qPCR) was performed using 10 ng of cDNA under the following amplification conditions: an initial 2Lmin denaturation at 95°C, followed by 40 cycles of 15 s at 95°C and 1 min at 60°C. Target mRNA expression was normalized to *Hypoxanthine phosphoribosyltransferase 1* (*HPRT*) as an endogenous control. Relative fold change was calculated using the 2^−double^ ^deltaCt^ method (Livak & Schmittgen, 2001).

### RNAScope

Fluorescent In Situ Hybridization was performed using the RNAscope Multiplex Fluorescence V2 Assay with pre-made RNAscope target probes C1–3 for human IL17B, TNF, and IL6 (Cat # 323100, Advanced Cell Diagnostics, Newark, CA). FFPE sections of human gastrinomas were baked and deparaffinized as previously described. RNAscope Multiplex fluorescence assay was performed according to the manufacturer’s kit instructions. Targets were visualized by developing the HRP signal with Akoya Opal 520, 570, 690 fluorophores according to the protocol using 1:750 dilution of each fluorophore (Cat #FP1488001, #FP1497001, #FP1501001, Akoya Biosciences, Marlborough, MA). Slides were then mounted using Prolong Gold anti-Fade mounting media with DAPI and imaged at 600X magnification using the Nikon Ti2-E inverted microscope system (Nikon, Tokyo, Japan).

### Statistical Analysis

The Benjamini-Hochberg multiple test correction was applied to all DSP analyses prior to testing for statistical significance. For samples with unequal variance, the nonparametric Mann- Whitney test was applied. For cases with equal variance, unpaired Student’s T-test with Tukey post-test was applied for comparisons involving two groups, while one-way ANOVA with Sidak post-test was used for comparisons involved three or more groups.

## Results

### Gastrinomas and non-functional NETs exhibit distinct neuro-immune protein signatures

Human duodenal gastrinomas express markers of neuroglial cells (Sundaresan *et al*. 2017) and neuroglial reprogramming was recently implicated in the development of pancreatic and pituitary NETs (Duan *et al*. 2022). To decipher whether GEP-NETs exhibit neuropathological features, we applied Nanostring Digital Spatial Profiling (DSP) technology to formalin-fixed paraffin-embedded (FFPE) tissue sections of three pancreatic gastrinomas (PGAST), two non-functioning pancreatic NETs (NF-PNET), and four duodenal gastrinomas (DGAST) (Table 1). We quantified the expression of a 40-plex human neuro-immune protein panel enriched with markers that are classically associated with Alzheimer’s and Parkinson’s diseases (Table 2). Consistent with previous reports (Rico *et al*. 2021, Elvis-Offiah *et al*. 2023), hematoxylin and eosin (H&E) stained images of DGASTs, PGASTs, and NF-PNETs showed the strongest stromal reaction in DGASTs (Figure 1A–C). Adjacent serial FFPE sections were stained with the Nanostring tumor morphology markers, pan-cytokeratin (PanCK) and CD45, labeling epithelium and immune cells, respectively. Using these markers, we selectively analyzed the expression of the neuro-immune protein panel in eighty 200- to 300-square micron regions of interest (ROIs) spanning tumor lesions, stroma, and adjacent normal epithelium including the Brunner’s glands (Figure 1D–F).

**Figure 1.**
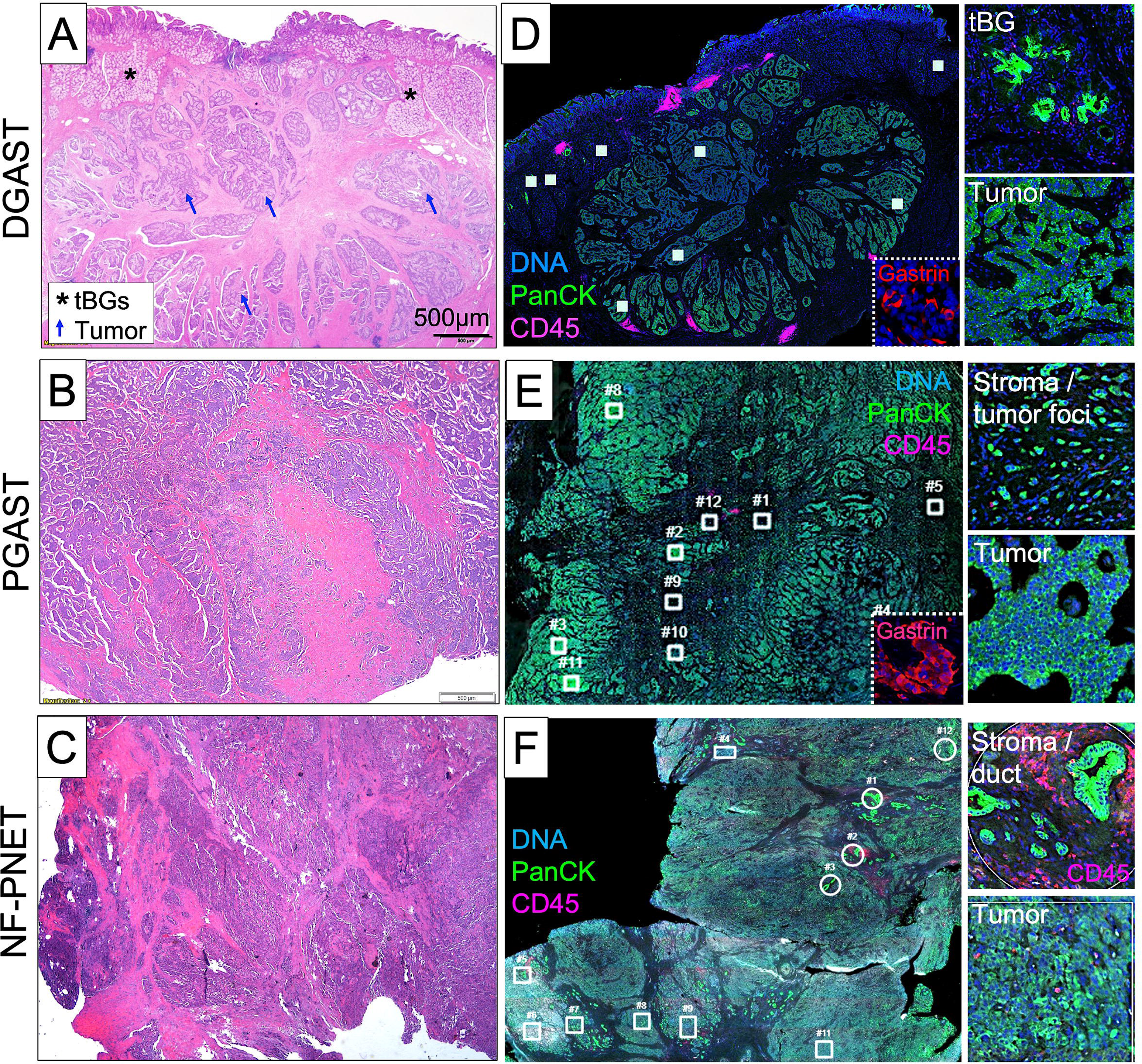
Digital spatial profiling of duodenal and pancreatic neuroendocrine tumors. *(A)* Hematoxylin and eosin-stained images of a duodenal gastrinoma (DGAST), *(B)* pancreatic gastrinoma (PGAST), and *(C)* non-functional pancreatic NET (NF-PNET). *(D)* Serial formalin- fixed paraffin-embedded sections of a DGAST stained with the GeoMx tumor morphology markers Pan cytokeratin (PanCK, green), CD45 (magenta), and SYTO13 (blue). Inset shows gastrin staining in red. White boxes and circles indicate regions of interest (ROIs) from which oligo-conjugated antibodies were collected for multi-plexed protein analysis. Right panel shows representative higher magnification images of the transitioning Brunner’s glands (tBGs) and tumor selected as regions of interest. *(E)* A PGAST and *(F)* NF-PNET stained with the GeoMx tumor morphology markers, with right panels showing representative ROIs of adjacent tumor stroma and tumor.

**Table 1.** Patient-derived duodenal and pancreatic NETs included in the discovery cohort for digital spatial profiling. The DSP discovery cohort included a total of 9 tumor-bearing patients and consisted of four duodenal gastrinomas (DGAST), three pancreatic gastrinomas (PGAST), and two non-functional PNETs (NF-PNETs). Known clinical and patient information including age, MEN1 status, and the presence of Zollinger-Ellison Syndrome (ZES), are also included. UNK denotes a patient with unknown details beyond the tumor diagnosis.

| Identification | Clinical diagnosis | Age | MEN1 syndrome | MEN1 mutation | ZES | Grade | Ki-67 | Mets | DSP |
| --- | --- | --- | --- | --- | --- | --- | --- | --- | --- |
| M2 | Duodenal gastrinoma | 57 | - | c.1621G>A | - | 1 | 2% | + | + |
| M4 | Duodenal gastrinoma | 52 | + | c.784-9G>A | + |  |  |  | + |
| M5 | Duodenal gastrinoma |  | UNK |  | UNK |  |  |  | + |
| M6, M6A | Duodenal gastrinoma |  | UNK |  | UNK |  |  |  | + |
| M8 | Non-functional PNET | 67 | - | c.703G>A | - | 2 | 7% | + | + |
| M9 | Non-functional PNET | 74 | - |  | - | 2 | 9% | + | + |
| M13 | Pancreatic gastrinoma |  | UNK |  | UNK |  |  |  | + |
| M14 | Pancreatic gastrinoma |  | UNK |  | UNK |  |  |  | + |
| M15 | Pancreatic gastrinoma |  | UNK |  | UNK |  |  |  | + |

**Table 2.** Protein targets and antibody controls included in Nanostring digital spatial profiling of human duodenal and pancreatic NETs. Digital spatial profiling was performed using the Nanostring GeoMx Neuro Cell Profiling Core target panel supplemented with additional neural- and immune-related protein targets included in the Alzheimer’s Disease and Parkinson’s Disease Pathology Modules. Histone H3, GAPDH, and S6 proteins were included as endogenous protein controls for normalizing target protein expression, while IgG controls were used for background signal subtraction.

| Controls | Neural Cell Profiling Core | Alzheimer's Disease Pathology Module | Parkinson's Disease Pathology Module |
| --- | --- | --- | --- |
| GAPDH | CD68 | Amyloid Precursor Protein | Alpha-synuclein |
| Histone H3 | HLA-DR | Amyloid-Beta 1-40 | ApoA-I |
| S6 | CD11b | Amyloid-Beta 1-42 | Calbindin |
| Rb IgG | MAP2 | APOE | FUS |
| Ms IgG | Ki-67 | P2RX7 | LRRK2 |
| Ms IgG2a | CD163 | Phospho-Tau (S404) | Park5 |
|  | Synaptophysin | Phospho-Tdp-43 (S409/S410) | Park7 |
|  | IBA1 | Tau | Phospho-Alpha-synuclein (S129) |
|  | GFAP | Tdp-43 | PINK1 |
|  | Myelin basic protein | Ubiquitin | Tyrosine Hydroxylase |
|  | NeuN |  |  |
|  | S100B |  |  |
|  | Olig2 |  |  |
|  | CD40 |  |  |
|  | CD45 |  |  |
|  | Neurofilament light |  |  |
|  | CD31 |  |  |
|  | CD39 |  |  |
|  | P2ry12 |  |  |
|  | TMEM119 |  |  |

Principal component analysis (PCA) was performed on tumor ROIs across DGASTs, PGASTs, and NF-PNETs to identify whether tumors exhibit distinct clustering based on their neuro-immune profiles (Figure 2A). Indeed, we observed clustering of the three tumor types with immune marker enrichment defining the first principal component (x-axis) and Ki-67 and neural protein expression defining the second principal component (y-axis) (Figure 2A). Protein expression analysis of the tumor ROIs showed remarkably increased expression of immune markers in PGASTs and NF-PNETs compared to the DGASTs (Figure 2B). PGASTs and NF- PNETs showed elevated expression of CD45 and the leukocyte-specific receptor CD11b (Figure 2C). Further, the pancreatic tumors exhibited distinct immune expression profiles, with PGASTs showing elevated expression of CD163 and CD39, marking myeloid cells and Foxp3^+^ regulatory T cells, respectively (Borsellino *et al*. 2007, Tremble *et al*. 2020). In comparison, NF-PNETs showed elevated CD68 and CD40 expression, primarily expressed by macrophages, dendritic cells, and B cells (Ma *et al*. 2009, Tremble *et al*. 2020). Consistent with the clinical pathology, NF-PNETs showed higher expression of Ki-67 expression compared to the other tumors in this cohort (Table 1 and Figure 2C). Immunohistochemical staining of the tumors confirmed these observations by showing reduced immune cell infiltrate and Ki-67 expression in DGASTs compared to the pancreatic NETs (Figure 2D). Therefore, the application of DSP to this cohort of GEP-NETs unveiled distinct immune profiles across tumor types and suggests fundamental differences in the neuro-immune cell composition of the tumor microenvironment.

**Figure 2.**
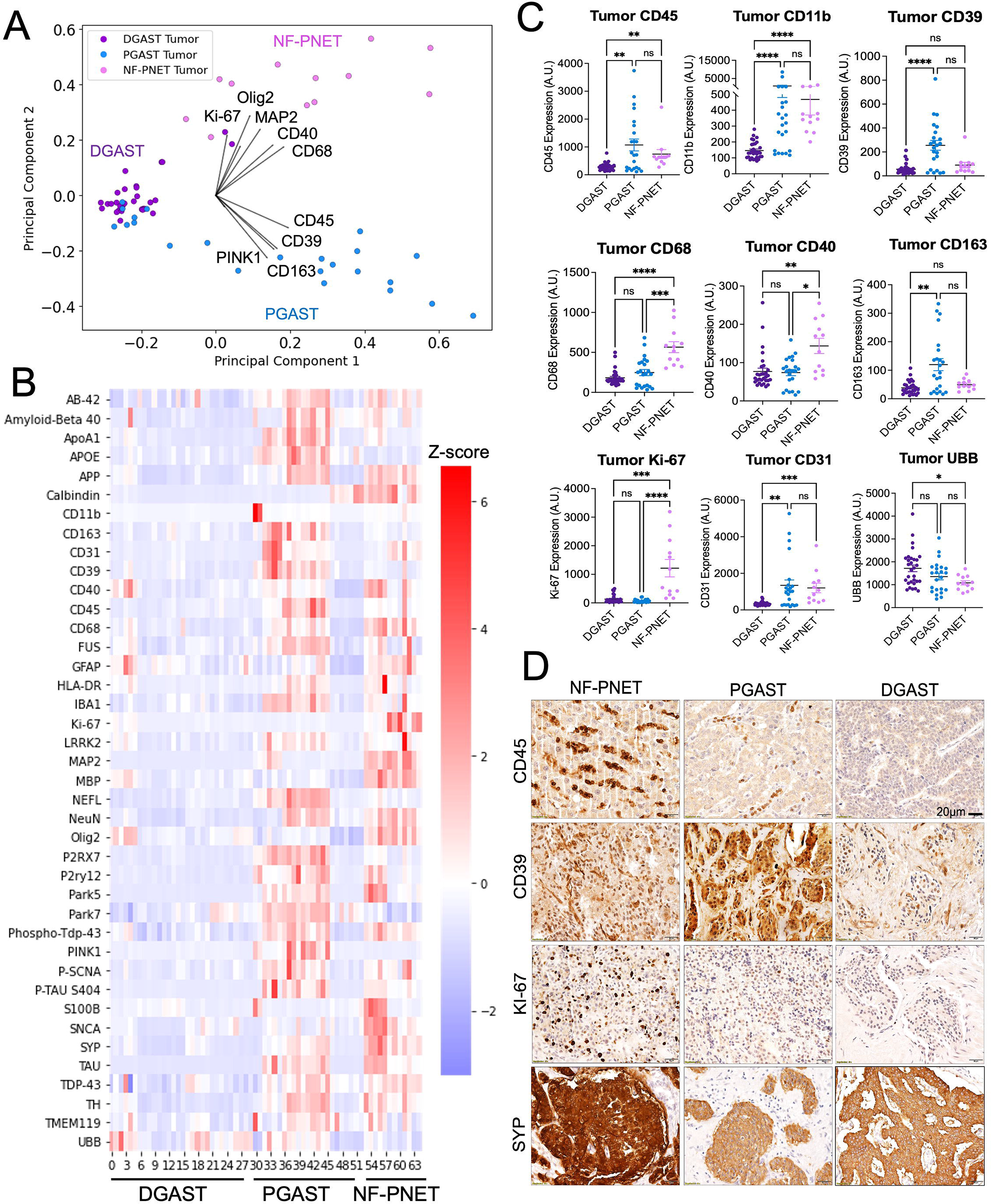
Gastrinomas and non-functional NETs exhibit distinct neuro-immune protein signatures. *(A)* Principal Component Analysis (PCA) plot of neuro-immune protein expression in tumor ROIs of DGASTs (n=3 patients, n=30 ROIs), PGASTs (n=3 patients, n=23 ROIs), and NF-PNETs (n=2 tumors, n=12 ROIs). Each dot represents a unique tumor ROI. *(B)* Heatmap showing relative expression (Z-score) of the 40 proteins analyzed in the DSP neuro-immune profiling panel. ROIs 0–29 indicate DGAST tumors, ROIs 30–52 indicate PGAST tumors, and ROIs 53–65 indicate NF-PNET tumors. *(C)* Tumor expression of select immune-related protein markers, in addition to the proliferative marker Ki-67, the endothelial marker CD31, and Ubiquitin (UBB). *(D)* Representative immunohistochemical (IHC) stained images showing the expression of CD45, CD39, Ki-67, and the neuroendocrine marker SYP in a DGAST, PGAST, and NF-PNET.

### GEP-NETs show strong intra- and inter-tumoral variation in neuro-immune protein expression

To better define the intra-tumoral abundance of these neuro-immune proteins, we compared their expression across normal adjacent tissue, tumor stroma, and tumor lesions. We further included a normal pancreas specimen (nPANC) and normal duodenum (nDUO) in our analysis for further comparison. PCA of the pancreas specimens separated nPANC and normal adjacent tissue from PGAST and NF-PNET stroma and tumor ROIs (Figure 3A and Supplementary Figure 1). The relative expression of neural and immune-related proteins defined these tissue classes along the second component (y-axis), with immune proteins upregulated in nPANC and normal adjacent tissues, and neural markers enriched in both the tumor stroma and tumor lesions (Figure 3A). As expected, tumor ROIs in both PGASTs and NF-PNETs strongly expressed the neuroendocrine marker SYP. However, PGASTs and NF-PNET tumor lesions showed differing expression levels of neuroglial markers, with PGASTs showing reduced expression of the glial protein S100B and increased expression of the Parkinson’s-associated protein, PTEN-induced kinase (PINK1). In contrast, NF-PNETs showed higher expression of proteins marking the neuroglial lineage, including MAP2 and OLIG2 (Figure 3B).

**Figure 3.**
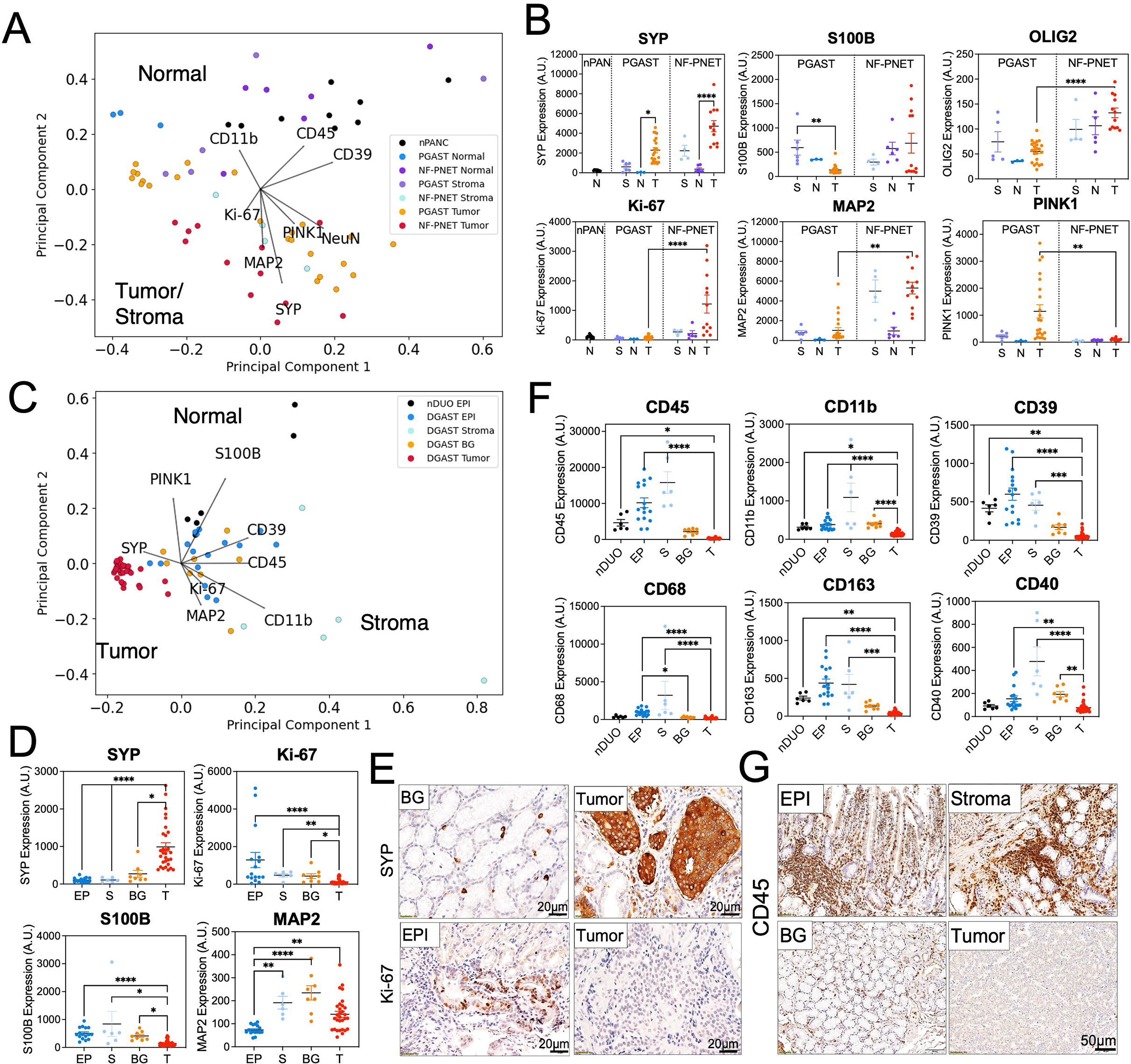
GEP-NETs show strong intra- and inter-tumoral variation in neuro-immune protein expression. *(A)* PCA plot showing the variation in DSP protein expression across pancreas tissues. ROIs were taken from a normal pancreas specimen for comparison (nPANC, n=9 ROIs from one patient). The three patient PGASTs were further divided into PGAST stroma (S, n=6 ROIs), PGAST normal adjacent tissue (N, n=3 ROIs), and PGAST tumor (T, n=23 ROIs). The two patient NF-PNETs were divided into NF-PNET stroma (n=4), NF-PNET normal adjacent tissue (n=6), and NF-PNET tumor (n=12). *(B)* Normalized protein expression of select markers across stroma, normal adjacent, and tumor ROIs of both PGASTs and NF-PNETs, respectively. *=*p* < 0.05, **=*p* < 0.01, ****=*p* < 0.0001 by one-way ANOVA with Tukey post-test. *(C)* PCA plot showing intra-tissue variation across the four patient DGASTs and a normal duodenum specimen (nDUO EPI, n=6 ROIs). DGASTs were subdivided into adjacent epithelium (EP, n=16 ROIs), stroma (S, n=6 ROIs), normal-appearing Brunner’s glands (BG, n=8 ROIs), and tumor (T, n=30 ROIs). *(D)* Normalized protein expression of select markers across tissue categories. *(E)* Representative images of immunohistochemical staining for SYP and Ki-67 showing appropriate expression across the tissue categories. *(F)* Normalized protein expression of immune-related proteins across tissue categories in the four patient DGASTs. *=*p* < 0.05, **=*p* < 0.01, ***=*p* < 0.001, ****=*p* < 0.0001 by one-way ANOVA with Tukey post-test. *(G)* Representative images of CD45 expression in the adjacent epithelium, normal-appearing BGs, stroma, and DGAST tumor lesion.

We next examined the intra-tissue variation of these proteins in our cohort of DGASTs by comparing expression levels across the nDUO specimen, adjacent epithelium, tumor stroma, adjacent Brunner’s glands, and tumor lesions. While the tumor ROIs clustered together strongly, the non-tumor compartments showed significant variation across the PCA (Figure 3C and Supplementary Figure 1). Similar to the pancreatic tumors, DGAST tumor ROIs showed significant enrichment in SYP expression but reduced expression of Ki-67 and S100B compared to adjacent tissues (Figure 3D). IHC staining of DGASTs and their adjacent tissues confirmed these expression patterns (Figure 3E). Most strikingly, DGAST tumor lesions showed strong exclusion of immune markers compared to the adjacent stroma and Brunner’s glands (Figure 3F and 3G). Taken together, GEP-NETs exhibit restricted immune profiles and upregulated neural protein expression compared to the surrounding tumor stroma and adjacent epithelium.

Furthermore, DGASTs and their associated Brunner’s glands show the highest extent of immune cell exclusion, consistent with a microenvironment that permits the reprogramming of these glands into preneoplastic lesions.

### Reduced immune infiltration and increased neural protein abundance in DGASTs is consistent in an expanded patient cohort

Given that the previous observations were made in a small patient population, we sought to extend these analyses to a broader cohort of GEP-NET-bearing patients with additional clinical annotation, including a diagnosis of MEN1 and Zollinger-Ellison Syndromes (ZES) (Table 3). We focused our subsequent investigation on DGASTs since these tumors are known to present with precursor lesions in the Brunner’s glands (Anlauf *et al*. 2005), thus marking them as an ideal model to study neoplastic development and tumor evolution. To investigate whether DGASTs upregulate their expression of neuroglial proteins during neoplastic development, we performed DSP on FFPE tumor sections from 12 patients, of which 11 were unique from the previous discovery cohort. We performed ROI selection on adjacent normal-appearing Brunner’s glands (nBG) and tumor lesions, in addition to transitioning Brunner’s glands (tBG) characterized by abnormal morphology, stromal thickening, and enrichment in PanCK expression. PCA and unbiased hierarchical clustering of 183 ROIs showed a coalescing phenotype with tBGs clustering between nBGs and tumor ROIs (Figure 4A and Supplementary Figure 2). As anticipated, SYP expression and enrichment in neural protein abundance strongly defined the tumor cluster (y-axis), while the nBG cluster (x-axis) was defined by elevated expression of immune-related markers (Figure 4A). Quantitative analysis of the three tissue classes showed a step wise increase in neural protein expression between tBGs and tumor lesions, and reduced expression of immune proteins, including markers of macrophage-like microglial cells (Jurga *et al*. 2020) (Figure 4B–D). IHC staining of the DGASTs confirmed increased expression of neuroendocrine and neural-related proteins SYP and alpha-synuclein (SNCA), in transitioning BGs and tumor lesions coincident with immune cell exclusion (Figure 4E). In summary, DGASTs and preneoplastic lesions display alterations in their neuro-immune protein profiles, suggesting that the immunologically cold microenvironment favors epithelial reprogramming and neoplastic transformation.

**Figure 4.**
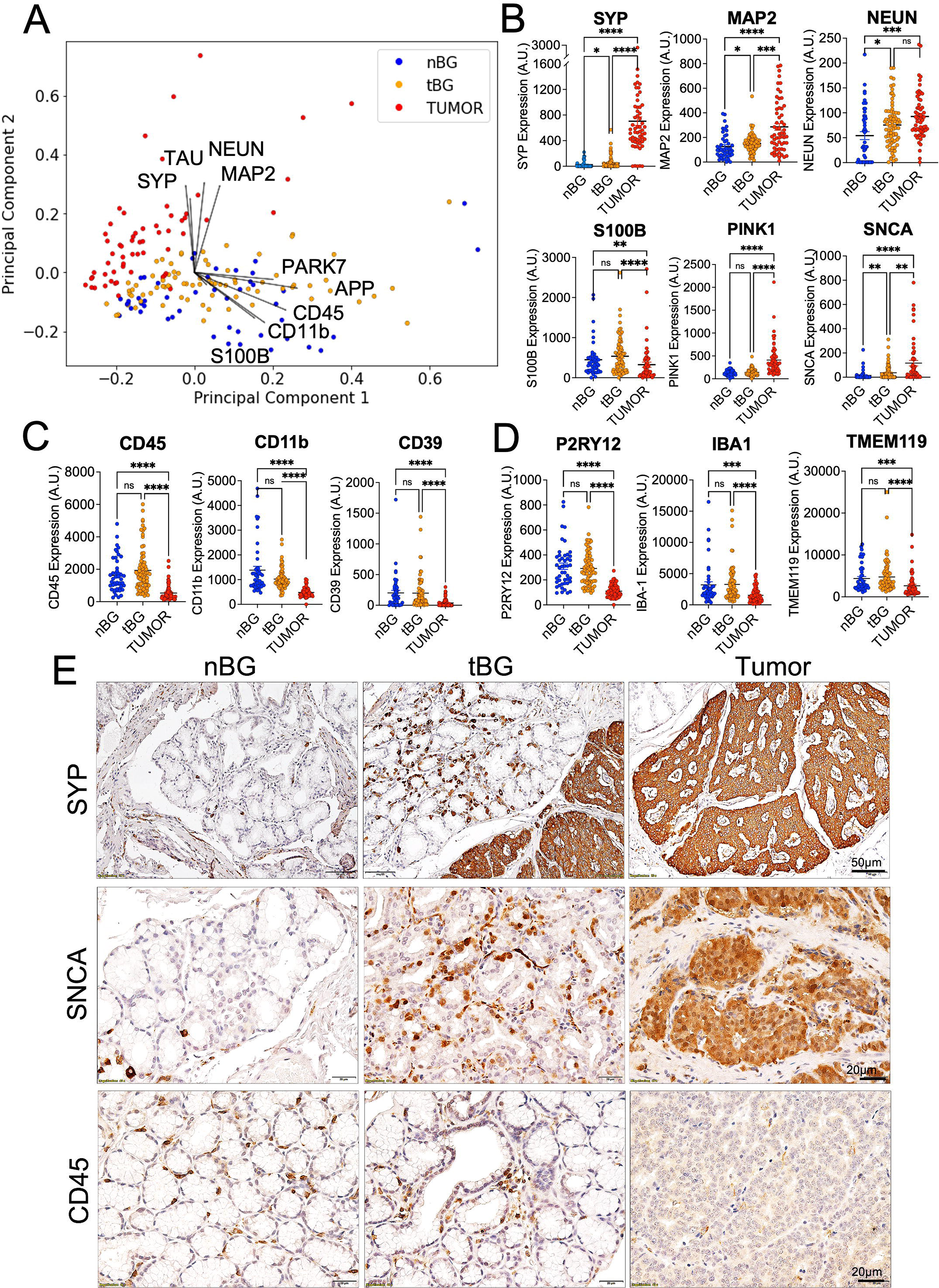
Reduced immune infiltration and increased neural protein abundance in DGASTs is consistent in an expanded patient cohort. *(A)* PCA plot showing the variation in DSP protein abundance across different tissue categories in DGASTs from 12 tumor-bearing patients. Regions analyzed include normal-appearing adjacent Brunner’s glands (nBG, n=47 ROIs), transitioning Brunner’s glands (tBG, n=76 ROIs), and tumor (n=60 ROIs). *(B)* Quantitation of select neuroendocrine and neuroglial proteins. *(C)* Quantitation of immune cell markers and *(D)* microglial-related proteins across the tissue types. *=*p* < 0.05, **=*p* < 0.01, ***=*p* < 0.001, ****=*p* < 0.0001 by one-way ANOVA with Tukey post-test. *(E)* Representative IHC stained imaged of nBGs, tBGs, and DGASTs showing expression of the neuroendocrine proteins SYP and SNCA, and the immune marker CD45.

**Table 3.**
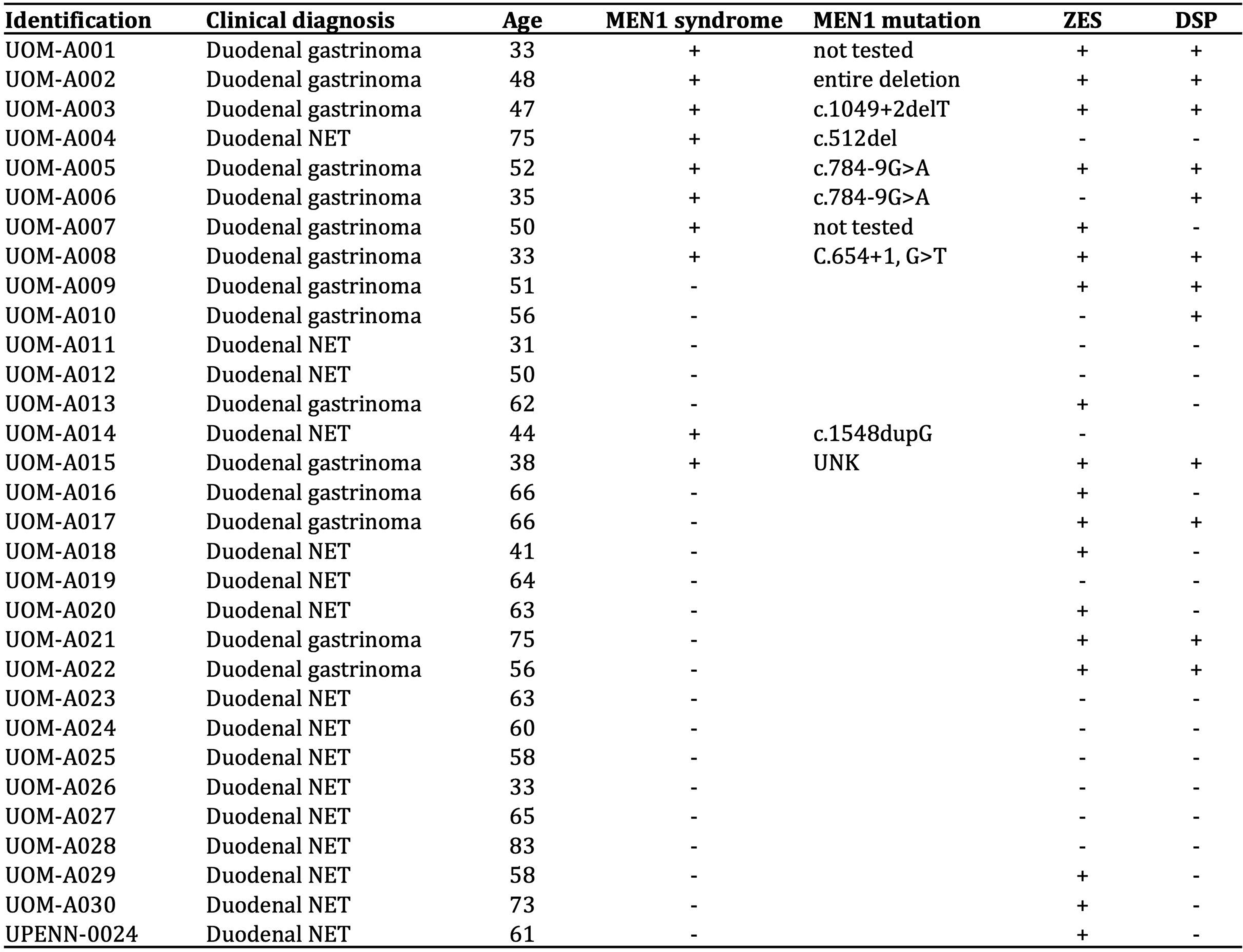
Patient demographics for duodenal gastrinomas and NETs included in the expanded digital spatial profiling analysis and validation studies. The expanded DSP analysis was performed on 12 DGAST-bearing patients, of which one patient had overlap in the previous discovery cohort. An additional 18 tumors, including DGASTs and non-gastrin secreting DNETs, were used for immunohistochemical staining validation. Relevant clinical information are also listed and include age, clinical MEN1 status, precise MEN1 mutations, and ZES status.

### MEN1 DGASTs show reduced immune infiltration and increased neuroglial protein expression compared to non-MEN1 tumors

Prior studies have indicated a role for menin, the protein product of the *MEN1* gene, in regulating cell fate specification leading to NET development (Sundaresan *et al*. 2017, Duan *et al*. 2022). Thus, we next investigated whether the neuro-immune profiles of DGASTs differed among patients with a MEN1 diagnosis. In our expanded cohort, 7/12 patients were clinically diagnosed with MEN1 syndrome (Table 3). We stratified the DSP results by tissue class and MEN1 syndrome and identified striking differences in the immune profiles of MEN1 DGASTs compared to non-MEN1 patients (Figure 5A and Supplementary Figure 2). Specifically, the tBGs of MEN1 patients showed significantly reduced expression of immune markers compared to the tBGs of non-MEN1 DGASTs, and this was confirmed upon IHC staining of the tumors (Figure 5B). Interestingly, the difference in immune expression grew less pronounced in tumor lesions from the two patient groups. Further, while SYP expression did not differ among the tissue classes in MEN1 versus non-MEN1 patients, MEN1 tumors showed significantly higher enrichment in neuroglial proteins compared to their non-MEN1 counterparts (Figure 5C). Lastly, non-MEN1 tBGs and tumor lesions exhibited significantly higher expression of Ki-67 and the endothelial marker CD31 (Figure 5D). These expression patterns were confirmed by IHC staining of DGAST lesions from both patient cohorts (Figure 5E). Thus, compared to non-MEN1 patients, the preneoplastic epithelium of MEN1 DGASTs exhibit a severely restricted immune microenvironment and MEN1 tumors show increased neuroglial differentiation and reduced proliferation.

**Figure 5.**
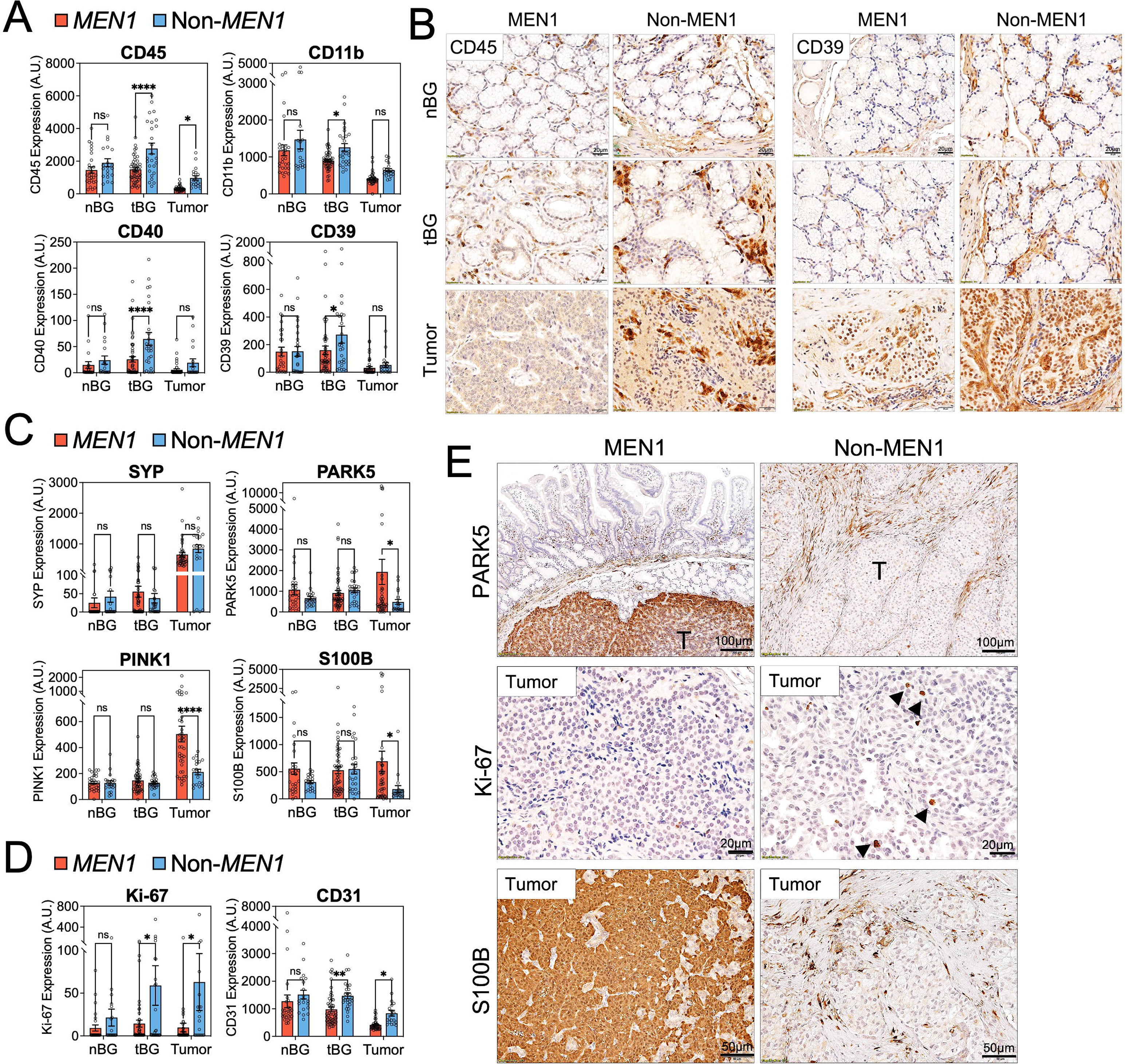
MEN1 DGASTs show reduced immune infiltration and increased neuroglial protein expression compared to non-MEN1 tumors. *(A)* Normalized expression of immune proteins across tissue categories in the 12 patient DGASTs stratified by MEN1 diagnosis. Protein expression was compared between normal-appearing adjacent Brunner’s glands (nBG MEN1 n=27, non-MEN1 n=20 ROIs), transitioning Brunner’s glands (tBG MEN1 n=51, non-MEN1 n=26 ROIs), and tumor (MEN1 n=40, non-MEN1 n=19 ROIs). *=*p* < 0.05, **=*p* < 0.01, ****=*p* < 0.0001 by two-way ANOVA with Benjamini-Hochberg multiple test correction. *(D)* Representative IHC stained images of CD45 and CD39 expression in nBG, tBG, and tumor lesions from MEN1 and non-MEN1 DGAST-bearing patients. *(C)* Normalized expression of neuroendocrine and neuroglial proteins and *(D)* Ki-67 and CD31 in the 12 patient DGASTs stratified by MEN1 status. *(E)* Representative IHC stained images of MEN1 and non-MEN1 DGASTs for Ki-67 and the neuroglial-associated proteins PARK5 and S100B. Arrows indicate nuclear Ki-67 expression.

### Tumor and stroma-derived IL-17B stimulate neuroendocrine differentiation downstream of NF-kB and STAT3 activation

Given that DGASTs express pro-inflammatory cytokines (Rico *et al*. 2021) and exhibit features of senescence, we sought to define whether DGASTs exhibit features of the senescence-associated secretory phenotype (SASP). We first confirmed that DGASTs and preneoplastic lesions in the expanded patient cohort express pro-inflammatory cytokines, including IL-17B, TNF11, and IL-6 (Figure 6A and B). Additionally, DGASTs were shown to express phospho- STAT3 (Tyr705) known to be activated downstream of these cytokines (Figure 6A). Consistent with SASP and low Ki-67 expression, DGASTs showed robust expression of the cyclin dependent kinase inhibitors and senescence markers p16^INK4^ and p21^Cip1^ (Figure 6C). We next evaluated the functional role of cytokine signaling in NET development using two primary human organoid lines that were generated from normal-appearing adjacent duodenum obtained during Whipple procedure. We focused our investigation on IL-17B as others have shown a role for IL-17 in altering cell fate specification and pro-neuronal differentiation (Lin *et al*. 2022, Zhang *et al*. 2023, Gomes *et al*. 2022). Moreover, of the six patient DGASTs examined by staining, four showed strong and diffuse IL-17B expression in SYP^+^ tumor cells, whereas the remaining 2/6 DGASTs and 3/3 PGASTs only showed sparse expression in the tumor stroma.

**Figure 6.**
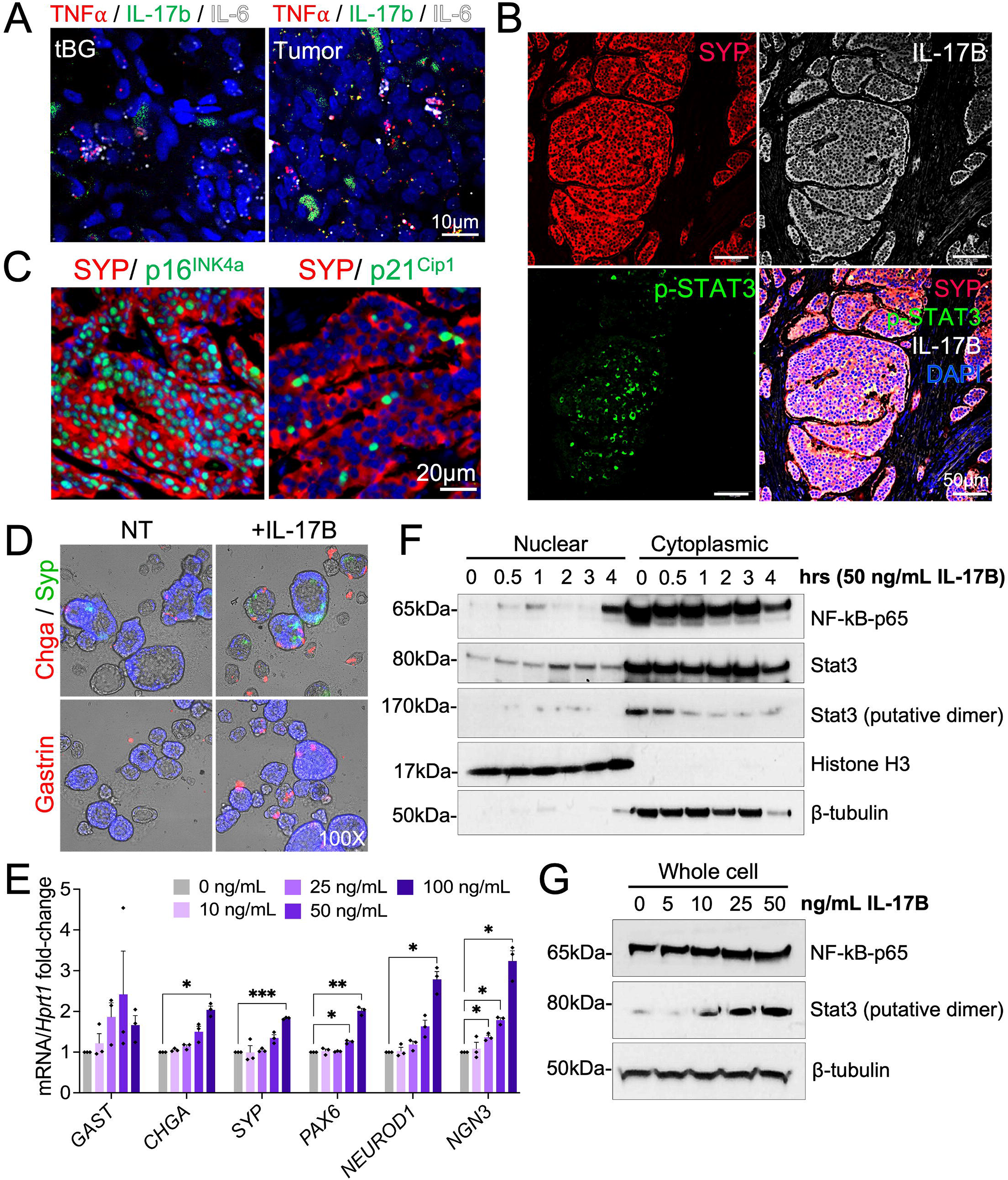
Tumor and stroma-derived IL-17B stimulate neuroendocrine differentiation downstream of NF-kB and STAT3 activation. *(A)* RNAscope of DGASTs and tBGs showing expression of *TNF-*lll, *IL-17B*, and *IL-6* in tumor lesions. *(B)* Representative immunofluorescent images of human DGAST tumors stained for IL-17B (white), phosho-STAT3 (Tyr705) (green), and SYP (red) protein expression. Four out of 6 patients showed strong, diffuse expression in the tumor. *(C)* Representative immunofluorescent images of DGASTs stained for the senescence markers p16^INK4^ and p21 (green) merged with SYP (red). *(D)* Immunofluorescent staining for the neuroendocrine markers SYP, CHGA, and gastrin in human duodenal organoids treated with 50 ng/mL IL-17B for 48 h. DAPI shown in blue. *(E)* Relative mRNA expression of neuroendocrine markers in human duodenal organoids treated with increasing concentrations of IL-17B for 48 h. n=3 experimental replicates with two patient organoid lines. *=*p* < 0.05, **=*p* < 0.01, ***=*p* < 0.001 by two-way ANOVA. *(F)* Expression of NF-kB and STAT3 proteins in nuclear and cytoplasmic lysates of human duodenal organoids treated with IL-17B for 0–4h. Histone H3 and β-tubulin shown as nuclear and cytoplasmic markers, respectively. *(G)* Expression of NF-kB and STAT3 in whole cell lysates of human duodenal organoids treated with increasing concentrations of IL-17B for 24 h.

Treatment of human duodenal organoids with IL-17B (50 ng/mL) stimulated the expression of chromogranin A (CHGA), SYP, and gastrin (Figure 6D). Similarly, IL-17B stimulated a dose-dependent increase in the expression of neuroendocrine transcripts including, *CHGA, PAX6, SYP, NEUROD1*, and *NGN3* (Figure 6E). We further showed the IL-17B activates the nuclear translocation of NF-kB/p65 and STAT3 proteins, suggesting that IL-17B acts through these pathways to alter the endocrine cell fate (Figure 6F and G).

Taken together, we show that DGASTs exhibit features of SASP, including upregulated expression of tumor-derived pro-inflammatory cytokines and cyclin-dependent kinase inhibitors. Moreover, we show that IL-17B exerts a functional effect on stimulating the neuroendocrine phenotype, consistent with prior reports showing that IL-17 alters cell fate commitment and neuronal differentiation. Thus, we summarize a potential mechanism of immune-autonomous IL- 17B signaling that contributes to the pathogenesis and unique molecular features of DGASTs (Figure 7).

**Figure 7.**
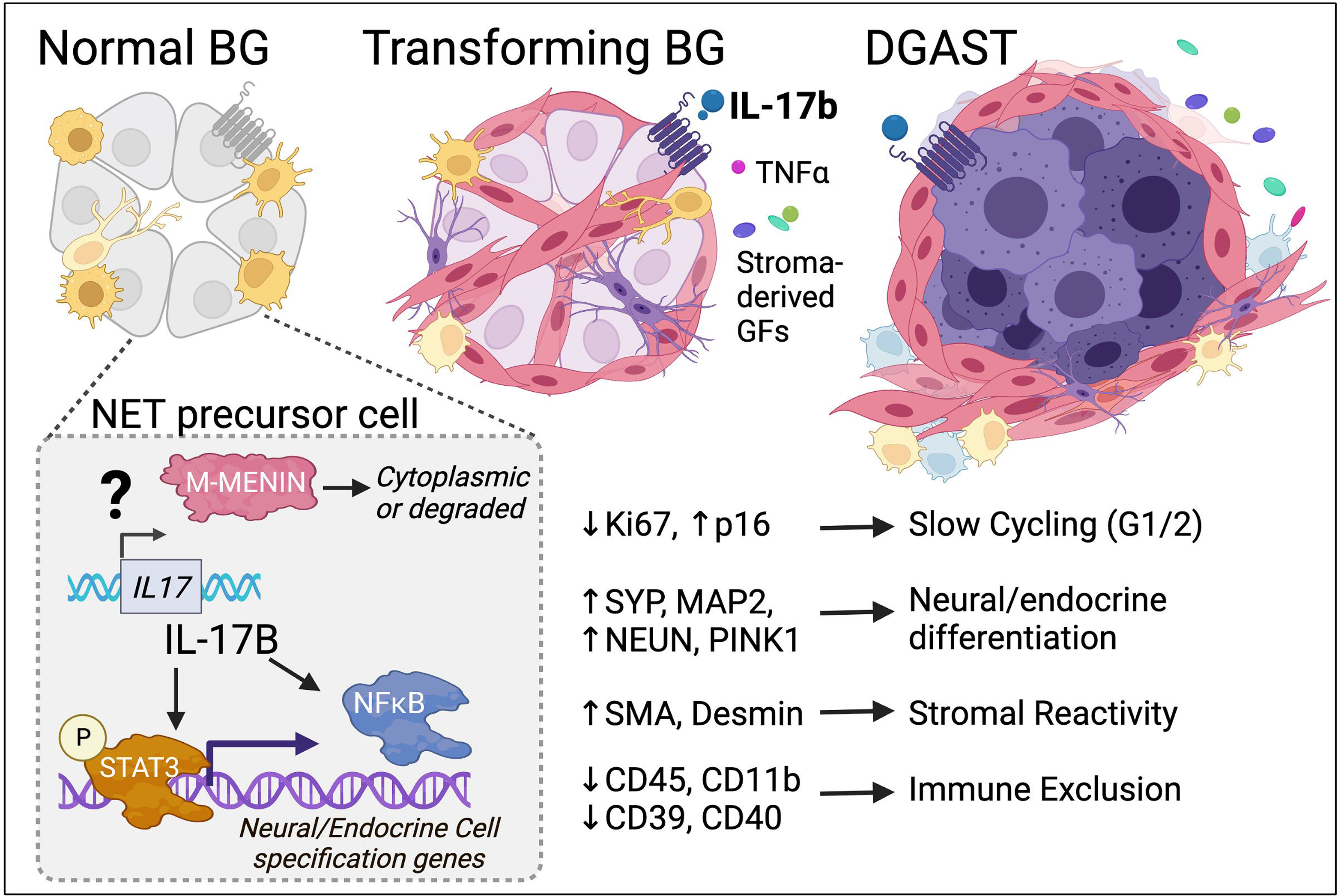
Potential mechanism of immune-autonomous IL-17b signaling in DGASTs. Under normal physiological conditions, the submucosal Brunner’s glands are comprised primarily of morphologically similar epithelial glands interspersed with immune cells. Enteroendocrine cells marked by chromogranin A and synaptophysin expression are relatively rare. Menin instability resulting from germline or somatic mutations may potentially derepress the expression of pro-inflammatory cytokines in stromal cells and the glandular epithelium. Upregulated IL-17B, a known pro-neuronal factor, acts on normal glandular tissue to induce neuroendocrine differentiation. The expression of IL-17B, TNF11, and IL-6 in DGASTs may be associated with the senescence associated secretory phenotype (SASP) that contributes to the classical features underlying well-differentiated G1/2 NETs, including slow proliferation, upregulated expression of CDKN2A/p16^INK4^, strong stromal reactivity leading to desmoplasia and finally, an immune excluded tumor microenvironment permissive to disease progression.

## Discussion

Past investigations into the molecular features of GEP-NETs are limited in their ability to spatially resolve the complex cellular interactions that shape the tumor microenvironment. To address this gap, we applied Nanostring Digital Spatial Profiling (DSP) technology to human GEP-NETs to profile the expression of a 40-plex human neuro-immune protein panel developed for neurological diseases, including Alzheimer’s and Parkinson’s diseases. Investigating the expression of these proteins is significant to understanding the disease process as human duodenal gastrinomas were previously shown to express markers of neuroglial cells and glial cells expressed gastrin in a genetic mouse model of gastric NET development (Sundaresan *et al*. 2017). Moreover, cellular reprogramming of neuroglial cells is implicated in the development of pancreatic and pituitary NETs in two glial-directed mouse models of the MEN1 syndrome (Duan *et al*. 2022). Thus, deciphering whether GEP-NETs exhibit shared neuropathological features may elucidate novel therapeutic approaches for targeting these malignancies.

Consistent with the previous reports, our study identified robust expression of neuroglial proteins in gastrinomas arising in the duodenum and pancreas. Distinct immune profiles further differentiated these tumors, with DGASTs and preneoplastic lesions exhibiting a microenvironment that strongly excludes immune cells. Similarly, a study examining the expression of PD-1, PD-L1, and T cell markers in small intestinal NETs (SI-NETs) and PNETs reported higher levels of T cell infiltrate in pancreatic tumors (da Silva *et al*. 2018). The immune-excluded environment shared across our cohort of DGASTs is consistent with other studies indicating low or absent PD-L1 and PD-1 expression in GEP-NETs (Sampedro-Nunez et al. 2018, Busse et al. 2020). Collectively, these studies help explain why immune checkpoint inhibition has seen limited success in GEP-NETs when administered as a monotherapy compared to other cancers with elevated tumor immune activity (Vijayvergia et al. 2020, Strosberg et al. 2020, Frumovitz *et al*. 2020).

However, these studies are more nuanced when other tumor features are considered. While well-differentiated GEP-NETs tend to show low objective response to adjuvant immunotherapies, patients with poorly differentiated G3 NETs and NECs experience improved outcomes, and higher response rates strongly correlated with the degree of immune infiltration in tumors (Bosch *et al*. 2019, Park *et al*. 2022). The high-grade non-functioning PNETs in our patient cohort exhibited the highest expression of immune markers, and this is consistent with other reports showing that G3 NETs and poorly differentiated NECs are associated with higher levels of tumor-infiltrating lymphocytes and lower recurrence-free survival (Ferrata et al. 2019, Bosch et al. 2019, Pucini et al. 2020, Busse et al. 2020). Surprisingly, MEN1-associated DGASTs in our cohort showed the lowest expression of both immune markers and immune cells in the adjacent transitioning Brunner’s glands. This suggests that immune exclusion in preneoplastic tissues and precursor lesions may promote a tumor permissive microenvironment that contributes to neuroendocrine differentiation and tumor progression (Figure 7). Thus, the decision to pursue adjuvant immunotherapy for GEP-NETs should be guided by additional factors, including the presence of MEN1 mutations, tumor differentiation status, and the site of tumor development.

Emerging studies have unveiled a role for the *MEN1* gene and its protein product menin, in regulating cytokine expression and driving cellular senescence (Kuwahara *et al*. 2014, Leng *et al*. 2018, Leng *at al.* 2023). Here, we hypothesized a role for cell autonomous cytokine signaling in tumor development. Expanding from our prior genomic analysis (Rico *et al*. 2021), we showed that DGASTs upregulate their expression of IL-17B independent of immune signaling, and this pro-neuronal cytokine plays a functional role in neuroendocrine reprogramming. Importantly, it remains unknown if menin regulates *IL17B* gene expression and whether *MEN1* mutations that destabilize menin expression derepress the *IL17B* promoter. Additional pull-down studies aimed at investigating these protein and chromatin interactions may illuminate potential avenues for therapeutic intervention in MEN1 GEP-NETs.

Finally, given that DGASTs exhibit features of the senescence-associated secretory phenotype, including cell cycle arrest (*e.g.* low Ki-67, high *CDKN2A*/p16^INK4^ expression), increased production of cytokines (e.g. IL-17B and TNF11), immune evasion, strong desmoplastic reaction, and neuronal-differentiation, future work to understand these mechanisms is strongly warranted. A specific role for IL-17B in mediating these events in DGASTs is emphasized by our observation that SYP^+^ tumor cells and preneoplastic lesions strongly express IL-17B (4/6 DGAST patients or 66%) compared to PGASTs that only showed stromal IL-17B expression. Thus, IL-17B may facilitate hallmarks unique to DGASTs, including stromal activation leading to immune exclusion (Chen *et al*. 2022). Reduced immune infiltrates in the transitioning Brunner’s glands and tumors occurred in the context of stromal thickening, suggesting that remodeling of the stroma underlies reprogramming of preneoplastic tissues into neoplastic lesions with neuroendocrine features.

In summary, spatial profiling technology enables a high through-put and quantitative approach that can spatially map numerous protein targets relevant to the disease process. Given their diverse heterogeneity and controversial cellular origins, GEP-NETs represent an ideal model for the application of this technology. Our study is among the first to profile the neuropathological features of DGASTs and shows that neuro-immune interactions play an important role in tissue reprogramming and disease progression.

## Author Contributions

SD designed and performed experimental studies, performed data analysis, and drafted the manuscript. TWS contributed to data visualization and analysis. BLW contributed to data analysis. HS contributed to immunohistochemical staining studies. TE collected and validated clinical samples. JLM conceptualized and designed the studies, edited the manuscript, and guaranteed funding.

## Data Availability Statement

Data are available upon reasonable request.

## Abbreviations

CHGA: Chromogranin A
DGAST: Duodenal Gastrinoma
DSP: Digital Spatial Profiling
FFPE: Formalin-fixed Paraffin-embedded
GEP-NET: Gastroenteropancreatic Neuroendocrine
Tumor IHC: Immunohistochemical Staining
MAP2: Microtubule-associated Protein
MEN1: Multiple Endocrine Neoplasia 1
nBG: Normal-appearing Brunner’s Glands
nDUO: Normal Human Duodenum
NEC: Neuroendocrine Carcinoma
NF-PNET: Non-functional PNET
nPANC: Normal Human Pancreas
PanCK: Pan Cyotkeratin
PCA: Principal Component Analysis
PGAST: Pancreatic Gastrinoma
PINK1: PTEN-induced Kinase
PNET: Pancreatic Neuroendocrine Tumor
ROI: Region of Interest
SASP: Senescence-associated Secretory Phenotype
SNCA: Alpha-synuclein
SI-NET: Small intestinal NET
SYP: Synaptophysin
tBG: Transitioning Brunner’s Glands
TME: Tumor Microenvironment
UBB: Ubiquitin

## Supplementary Data Legends

**Supplementary Table 1. Reagents for sub-culturing human duodenal organoids.**

Supplementary Figure 1. PGASTs, NF-PNETs, and DGASTs show strong intra-tissue and inter-tumor variation in the expression of neuro-immune protein markers. *(A)* Heatmap showing the expression of the Nanostring 40-plex protein panel across regions of interest selected from normal pancreas (nPANC) (n=1 patient), PGAST (n=3 patients), and NF-PNETs (n=2 patients). A total of n=9 ROIs were analyzed in the nPANC specimen. Tumor tissues were further divided into PGAST stroma (S, n=6 ROIs), PGAST normal adjacent tissue (N, n=3), PGAST tumor (T, n=23), NF-PNET stroma (n=4), NF-PNET normal adjacent tissue (n=6), and NF-PNET tumor (n=12). *(B)* Heatmap showing the expression of the 40-plex protein panel in the epithelium of a normal duodenum specimen (nDUO EP, n=1 patient, n=6 ROIs), compared to epithelium (EP, n=16), stroma (S, n=6), Brunner’s glands (BGs, n=8), and tumor (T, n=30) ROIs from 4 DGAST-bearing patients.

**Supplementary Figure 2. Unsupervised hierarchical clustering of DGASTs shows progressive changes in the neuro-immune phenotype during tumorigenesis.** Unbiased hierarchical clustering of DSP protein expression was performed across the 12 DGAST-bearing patients. For each specimen, the ROI was assigned to normal-appearing Brunner’s glands (nBGs, n=47 ROIs) transitioning Brunner’s glands (tBGs, n=76 ROIs), and tumor (T, n=60 ROIs) and denoted by color along the top y-axis. ROIs from a single patient are denoted by color along the bottom y-axis of the heatmap. Patients were further stratified by MEN1 status to identify differing expression patterns.

## References

Anlauf M, Perren A, Meyer CL, Schmid S, Saremaslani P, Kruse ML, Weihe E, Komminoth P, Heitz PU, Klöppel G. 2005. Precursor lesions in patients with multiple endocrine neoplasia type 1-associated duodenal gastrinomas. Gastroenterology, 128(5), 1187–1198.

Borsellino G, Kleinewietfeld M, Di Mitri D, Sternjak A, Diamantini A, Giometto R, Höpner S, Centonze D, Bernardi G, Dell’Acqua ML, Rossini PM, Battistini L, Rötzschke O, Falk K. 2007. Expression of ectonucleotidase CD39 by Foxp3+ Treg cells: hydrolysis of extracellular ATP and immune suppression. Blood, 110(4), 1225–1232.

Bösch F, Brüwer K, Altendorf-Hofmann A, Auernhammer CJ, Spitzweg C, Westphalen CB, Boeck S, Schubert-Fritschle G, Werner J, Heinemann V, Kirchner T, Angele M, Knösel T. 2019. Immune checkpoint markers in gastroenteropancreatic neuroendocrine neoplasia. Endocrine-related cancer, 26(3), 293–301.

Brady L, Kriner M, Coleman I, Morrissey C, Roudier M, True LD, Gulati R, Plymate SR, Zhou Z, Birditt B, Meredith R, Geiss G, Hoang M, Beechem J, Nelson PS. 2021. Inter- and intra-tumor heterogeneity of metastatic prostate cancer determined by digital spatial gene expression profiling. Nature communications, 12(1), 1426.

Busse A, Mochmann LH, Spenke C, Arsenic R, Briest F, Jöhrens K, Lammert H, Sipos B, Kühl AA, Wirtz R, Pavel M, Hummel M, Kaemmerer D, Baum RP, Grabowski P. 2020. Immunoprofiling in Neuroendocrine Neoplasms Unveil Immunosuppressive Microenvironment. Cancers, 12(11), 3448.

Carter JM, Chumsri S, Hinerfeld DA, Ma Y, Wang X, Zahrieh D, Hillman DW, Tenner KS, Kachergus JM, Brauer HA, Warren SE, Henderson D, Shi J, Liu Y, Joensuu H, Lindman H, Leon-Ferre RA, Boughey JC, Liu MC, Ingle JN, Kalari KR, Couch FJ, Knutson KL, Goetz MP, Perez EA, Thompson EA. 2023. Distinct spatial immune microlandscapes are independently associated with outcomes in triple-negative breast cancer. Nature communications, 14(1), 2215

da Silva A, Bowden M, Zhang S, Masugi Y, Thorner AR, Herbert ZT, Zhou CW, Brais L, Chan JA, Hodi FS, Rodig S, Ogino S, Kulke MH. 2018. Characterization of the Neuroendocrine Tumor Immune Microenvironment. Pancreas, 47(9), 1123–1129.

Duan S, Sawyer TW, Sontz RA, Wieland BA, Diaz AF, Merchant JL. 2022. GFAP- directed Inactivation of Men1 Exploits Glial Cell Plasticity in Favor of Neuroendocrine Reprogramming. Cellular and molecular gastroenterology and hepatology, 14(5), 1025– 1051.

Ferrata M, Schad A, Zimmer S, Musholt TJ, Bahr K, Kuenzel J, Becker S, Springer E, Roth W, Weber MM, Fottner C. 2019. PD-L1 Expression and Immune Cell Infiltration in Gastroenteropancreatic (GEP) and Non-GEP Neuroendocrine Neoplasms With High Proliferative Activity. Frontiers in oncology, 9, 343.

Frumovitz M, Westin SN, Salvo G, Zarifa A, Xu M, Yap TA, Rodon AJ, Karp DD, Abonofal A, Jazaeri AA, Naing A. 2020. Phase II study of pembrolizumab efficacy and safety in women with recurrent small cell neuroendocrine carcinoma of the lower genital tract. Gynecologic oncology, 158(3), 570–575.

Gomes AKS, Dantas RM, Yokota BY, Silva ALTE, Griesi-Oliveira K, Passos-Bueno MR, Sertié AL. 2022. Interleukin-17a Induces Neuronal Differentiation of Induced- Pluripotent Stem Cell-Derived Neural Progenitors From Autistic and Control Subjects. Frontiers in neuroscience, 16, 828646.

How-Kit A, Dejeux E, Dousset B, Renault V, Baudry M, Terris B, Tost J. 2015. DNA methylation profiles distinguish different subtypes of gastroenteropancreatic neuroendocrine tumors. Epigenomics, 7(8), 1245–1258.

Jensen RT, & Norton JA. 2017. Treatment of Pancreatic Neuroendocrine Tumors in Multiple Endocrine Neoplasia Type 1: Some Clarity But Continued Controversy. Pancreas, 46(5), 589–594.

Jiao Y, Shi C, Edil BH, de Wilde RF, Klimstra DS, Maitra A, Schulick RD, Tang LH, Wolfgang CL, Choti MA, Velculescu VE, Diaz LA Jr, Vogelstein B, Kinzler KW, Hruban RH, Papadopoulos N. 2011. DAXX/ATRX, MEN1, and mTOR pathway genes are frequently altered in pancreatic neuroendocrine tumors. Science (New York, N.Y.), 331(6021), 1199–1203.

Jurga AM, Paleczna M, Kuter KZ. 2020. Overview of General and Discriminating Markers of Differential Microglia Phenotypes. Frontiers in cellular neuroscience, 14, 198.

Kawasaki K, Fujii M, & Sato T. 2018. Gastroenteropancreatic neuroendocrine neoplasms: genes, therapies and models. Disease models & mechanisms, 11(2), dmm029595.

Kim Y, Danaher P, Cimino PJ, Hurth K, Warren S, Glod J, Beechem JM, Zada G, McEachron TA. 2023. Highly Multiplexed Spatially Resolved Proteomic and Transcriptional Profiling of the Glioblastoma Microenvironment Using Archived Formalin-Fixed Paraffin-Embedded Specimens. Modern pathology : an official journal of the United States and Canadian Academy of Pathology, Inc, 36(1), 100034.

Kuwahara M, Suzuki J, Tofukuji S, Yamada T, Kanoh M, Matsumoto A, Maruyama S, Kometani K, Kurosaki T, Ohara O, Nakayama T, Yamashita M. 2014. The Menin-Bach2 axis is critical for regulating CD4 T-cell senescence and cytokine homeostasis. Nature communications, 5, 3555.

Leng L, Zhuang K, Liu Z, Huang C, Gao Y, Chen G, Lin H, Hu Y, Wu D, Shi M, Xie W, Sun H, Shao Z, Li H, Zhang K, Mo W, Huang TY, Xue M, Yuan Z, Zhang X, Bu G, Xu H, Xu Q, Zhang J. 2018. Menin Deficiency Leads to Depressive-like Behaviors in Mice by Modulating Astrocyte-Mediated Neuroinflammation. Neuron, 100(3), 551–563.e7.

Leng L, Yuan Z, Su X, Chen Z, Yang S, Chen M, Zhuang K, Lin H, Sun H, Li H, Xue M, Xu J, Yan J, Chen Z, Yuan T, Zhang J. 2023. Hypothalamic Menin regulates systemic aging and cognitive decline. PLoS biology, 21(3), e3002033.

Lin X, Gaudino SJ, Jang KK, Bahadur T, Singh A, Banerjee A, Beaupre M, Chu T, Wong HT, Kim CK, Kempen C, Axelrad J, Huang H, Khalid S, Shah V, Eskiocak O, Parks OB, Berisha A, McAleer JP, Good M, Hoshino M, Blumberg R, Bialkowska AB, Gaffen SL, Kolls JK, Yang VW, Beyaz S, Cadwell K, Kumar P. 2022. IL-17RA- signaling in Lgr5^+^ intestinal stem cells induces expression of transcription factor ATOH1 to promote secretory cell lineage commitment. Immunity, 55(2), 237–253.e8.

Livak KJ, Schmittgen TD. 2001. Analysis of relative gene expression data using real-time quantitative PCR and the 2(-Delta Delta C(T)) Method. Methods. 25, 402–8.

Lou X, Gao H, Xu X, Ye Z, Zhang W, Wang F, Chen J, Zhang Y, Chen X, Qin Y, Yu X, Ji S. 2022. The Interplay of Four Main Pathways Recomposes Immune Landscape in Primary and Metastatic Gastroenteropancreatic Neuroendocrine Tumors. Frontiers in oncology, 12, 808448.

Ma DY, Clark EA. 2009. The role of CD40 and CD154/CD40L in dendritic cells. Seminars in immunology, 21(5), 265–272.

Melone V, Salvati A, Palumbo D, Giurato G, Nassa G, Rizzo F, Palo L, Giordano A, Incoronato M, Vitale M, Mian C, Di Biase I, Cristiano S, Narciso V, Cantile M, Di Mauro A, Tatangelo F, Tafuto S, Modica R, Pivonello C, Salvatore M, Colao A, Weisz A, Tarallo R. 2022. Identification of functional pathways and molecular signatures in neuroendocrine neoplasms by multi-omics analysis. Journal of translational medicine, 20(1), 306.

Panarelli N, Tyryshkin K, Wong JJM, Majewski A, Yang X, Scognamiglio T, Kim MK, Bogardus K, Tuschl T, Chen YT, Renwick N. 2019. Evaluating gastroenteropancreatic neuroendocrine tumors through microRNA sequencing. Endocrine-related cancer, 26(1), 47–57.

Puccini A, Poorman K, Salem ME, Soldato D, Seeber A, Goldberg RM, Shields AF, Xiu J, Battaglin F, Berger MD, Tokunaga R, Naseem M, Barzi A, Iqbal S, Zhang W, Soni S, Hwang JJ, Philip PA, Sciallero S, Korn WM, Marshall JL, Lenz HJ. 2020. Comprehensive Genomic Profiling of Gastroenteropancreatic Neuroendocrine Neoplasms (GEP-NENs). Clinical cancer research : an official journal of the American Association for Cancer Research, 26(22), 5943–5951.

Rico K, Duan S, Pandey RL, Chen Y, Chakrabarti JT, Starr J, Zavros Y, Else T, Katona B W, Metz DC, & Merchant JL. 2021. Genome analysis identifies differences in the transcriptional targets of duodenal versus pancreatic neuroendocrine tumours. BMJ open gastroenterology, 8(1), e000765.

Sampedro-Núñez M, Serrano-Somavilla A, Adrados M, Cameselle-Teijeiro JM, Blanco- Carrera C, Cabezas-Agricola JM, Martínez-Hernández R, Martín-Pérez E, Muñoz de Nova JL, Díaz JÁ, García-Centeno R, Caneiro-Gómez J, Abdulkader I, González-Amaro R, Marazuela M. 2018. Analysis of expression of the PD-1/PD-L1 immune checkpoint system and its prognostic impact in gastroenteropancreatic neuroendocrine tumors. Scientific reports, 8(1), 17812.

Scarpa A, Chang DK, Nones K, Corbo V, Patch AM, Bailey P, Lawlor RT, Johns AL, Miller DK, Mafficini A, Rusev B, Scardoni M, Antonello D, Barbi S, Sikora KO, Cingarlini S, Vicentini C, McKay S, Quinn MC, Bruxner TJ, Christ AN, Harliwong I, Idrisoglu S, McLean S, Nourse C, Nourbakhsh E, Wilson PJ, Anderson MJ, Fink JL, Newell F, Waddell N, Holmes O, Kazakoff SH, Leonard C, Wood S, Xu Q, Nagaraj SH, Amato E, Dalai I, Bersani S, Cataldo I, Dei Tos AP, Capelli P, Davì MV, Landoni L, Malpaga A, Miotto M, Whitehall VL, Leggett BA, Harris JL, Harris J, Jones MD, Humphris J, Chantrill LA, Chin V, Nagrial AM, Pajic M, Scarlett CJ, Pinho A, Rooman I, Toon C, Wu J, Pinese M, Cowley M, Barbour A, Mawson A, Humphrey ES, Colvin EK, Chou A, Lovell JA, Jamieson NB, Duthie F, Gingras MC, Fisher WE, Dagg RA, Lau LM, Lee M, Pickett HA, Reddel RR, Samra JS, Kench JG, Merrett ND, Epari K, Nguyen NQ, Zeps N, Falconi M, Simbolo M, Butturini G, Van Buren G, Partelli S, Fassan M; Australian Pancreatic Cancer Genome Initiative; Khanna KK, Gill AJ, Wheeler DA, Gibbs RA, Musgrove EA, Bassi C, Tortora G, Pederzoli P, Pearson JV, Waddell N, Biankin AV, Grimmond SM. 2017. Whole-genome landscape of pancreatic neuroendocrine tumours. Nature, 543(7643), 65–71.

Sokal RR, Michener CD. 1958. A statistical method for evaluating systematic relationships. University of Kansas Science Bulletin. 38, 1409–1438.

Stabile BE, Morrow DJ, & Passaro E Jr. 1984. The gastrinoma triangle: operative implications. American journal of surgery, 147(1), 25–31.

Strosberg J, Mizuno N, Doi T, Grande E, Delord JP, Shapira-Frommer R, Bergsland E, Shah M, Fakih M, Takahashi S, Piha-Paul SA, O’Neil B, Thomas S, Lolkema MP, Chen M, Ibrahim N, Norwood K, Hadoux J. 2020. Efficacy and Safety of Pembrolizumab in Previously Treated Advanced Neuroendocrine Tumors: Results From the Phase II KEYNOTE-158 Study. Clinical cancer research : an official journal of the American Association for Cancer Research, 26(9), 2124–2130.

Sundaresan S, Meininger CA, Kang AJ, Photenhauer AL, Hayes MM, Sahoo N, Grembecka J, Cierpicki T, Ding L, Giordano TJ, Else T, Madrigal DJ, Low MJ, Campbell F, Baker AM, Xu H, Wright NA, Merchant JL. 2017. Gastrin Induces Nuclear Export and Proteasome Degradation of Menin in Enteric Glial Cells. Gastroenterology, 153(6), 1555–1567.e15.

Tremble LF, McCabe M, Walker SP, McCarthy S, Tynan RF, Beecher S, Werner R, Clover AJP, Power XDG, Forde PF, Heffron CCBB. 2020. Differential association of CD68^+^ and CD163^+^ macrophages with macrophage enzymes, whole tumour gene expression and overall survival in advanced melanoma. British journal of cancer, 123(10), 1553–1561.

Vijayvergia N, Dasari A, Deng M, Litwin S, Al-Toubah T, Alpaugh RK, Dotan E, Hall MJ, Ross NM, Runyen MM, Denlinger CS, Halperin DM, Cohen SJ, Engstrom PF, Strosberg JR. 2020. Pembrolizumab monotherapy in patients with previously treated metastatic high-grade neuroendocrine neoplasms: joint analysis of two prospective, non-randomised trials. British journal of cancer, 122(9), 1309–1314.

Wu H, Yu Z, Liu Y, Guo L, Teng L, Guo L, Liang L, Wang J, Gao J, Li R, Yang L, Nie X, Su D, Liang Z. 2022. Genomic characterization reveals distinct mutation landscapes and therapeutic implications in neuroendocrine carcinomas of the gastrointestinal tract. *Cancer communications (London*, England), 42(12), 1367–1386.

Yang B, Li X, Zhang W, Fan J, Zhou Y, Li W, Yin J, Yang X, Guo E, Li X, Fu Y, Liu S, Hu D, Qin X, Dou Y, Xiao R, Lu F, Wang Z, Qin T, Wang W, Zhang Q, Li S, Ma D, Mills GB, Chen G, Sun C. 2022. Spatial heterogeneity of infiltrating T cells in high-grade serous ovarian cancer revealed by multi-omics analysis. *Cell reports*. Medicine, 3(12), 100856.

Zhang Y, Zoltan M, Riquelme E, Xu H, Sahin I, Castro-Pando S, Montiel MF, Chang K, Jiang Z, Ling J, Gupta S, Horne W, Pruski M, Wang H, Sun SC, Lozano G, Chiao P, Maitra A, Leach SD, Kolls JK, Vilar E, Wang TC, Bailey JM, McAllister F. 2018. Immune Cell Production of Interleukin 17 Induces Stem Cell Features of Pancreatic Intraepithelial Neoplasia Cells. Gastroenterology, 155(1), 210–223.e3.

Chen X, Zhao J, Herjan T, Hong L, Liao Y, Liu C, Vasu K, Wang H, Thompson A, Fox PL, Gastman BR, Li X, Li X. 2022. IL-17-induced HIF1α drives resistance to anti-PD-L1 via fibroblast-mediated immune exclusion. The Journal of experimental medicine, 219(6), e20210693.

