## Supplementary figures and images for "Spatial profiling of neuro-immune interactions in gastroenteropancreatic NETs"

### Supplementary Figure 1

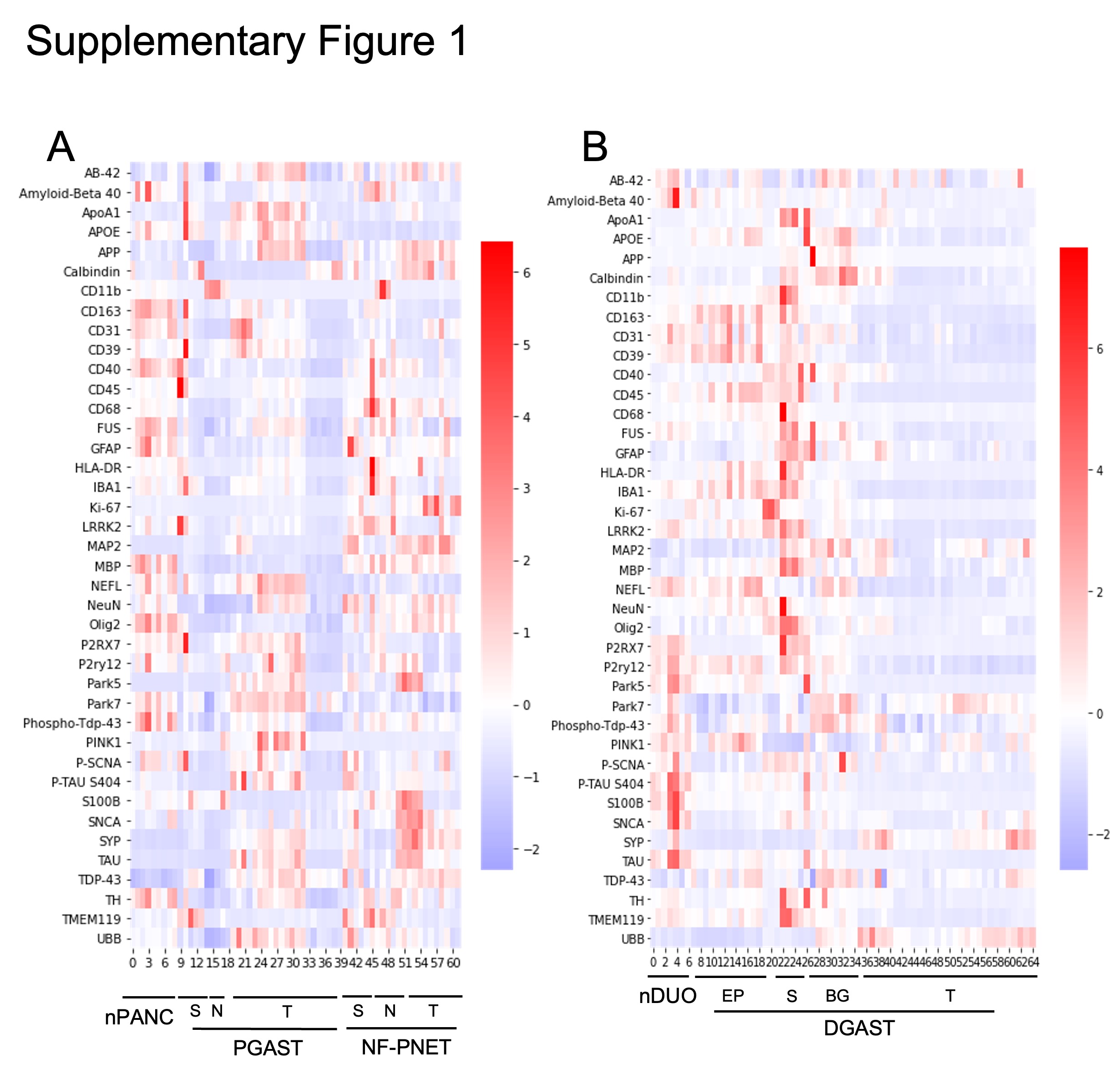

### Supplementary Figure 2

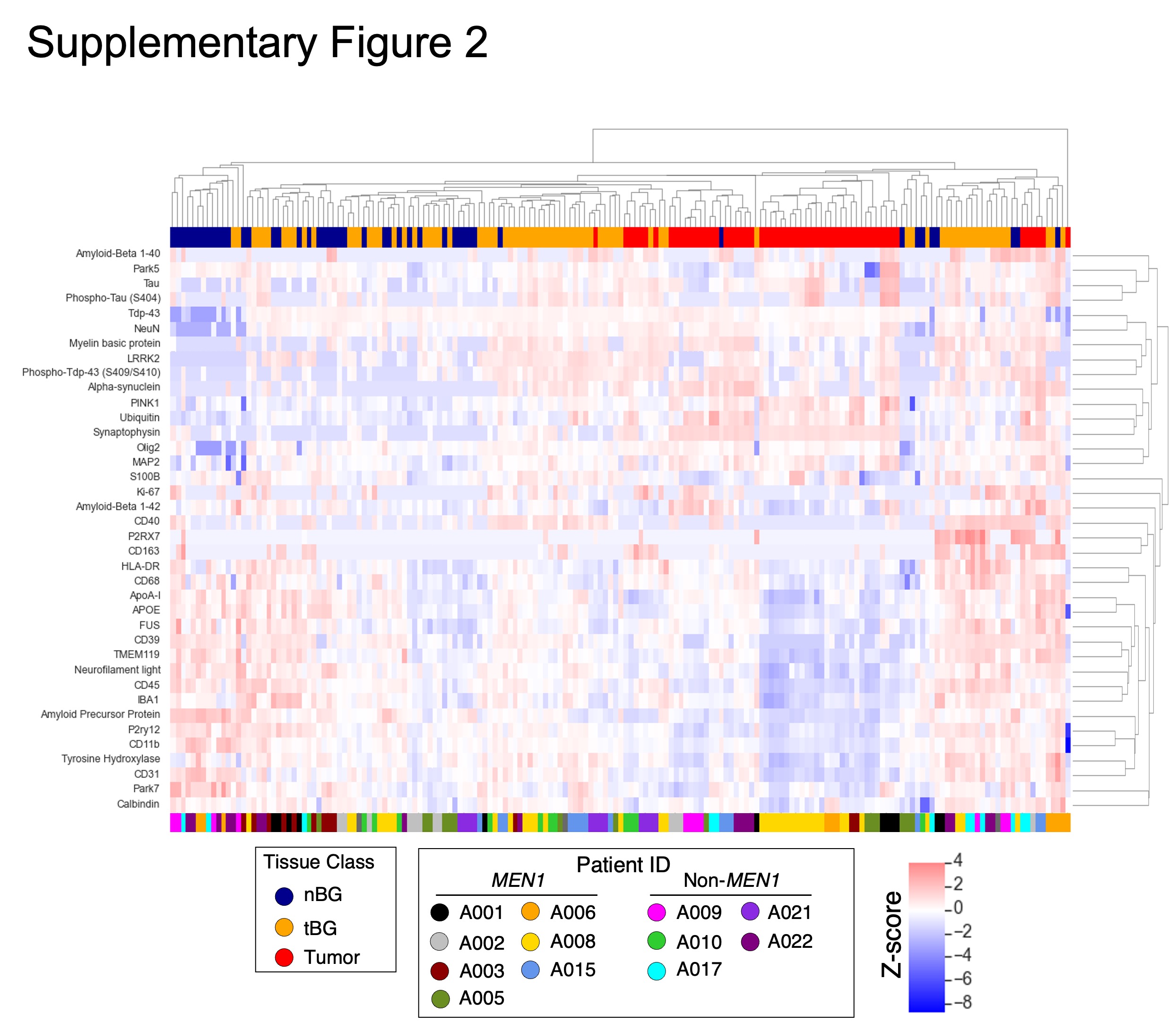

### Supplementary Table 1

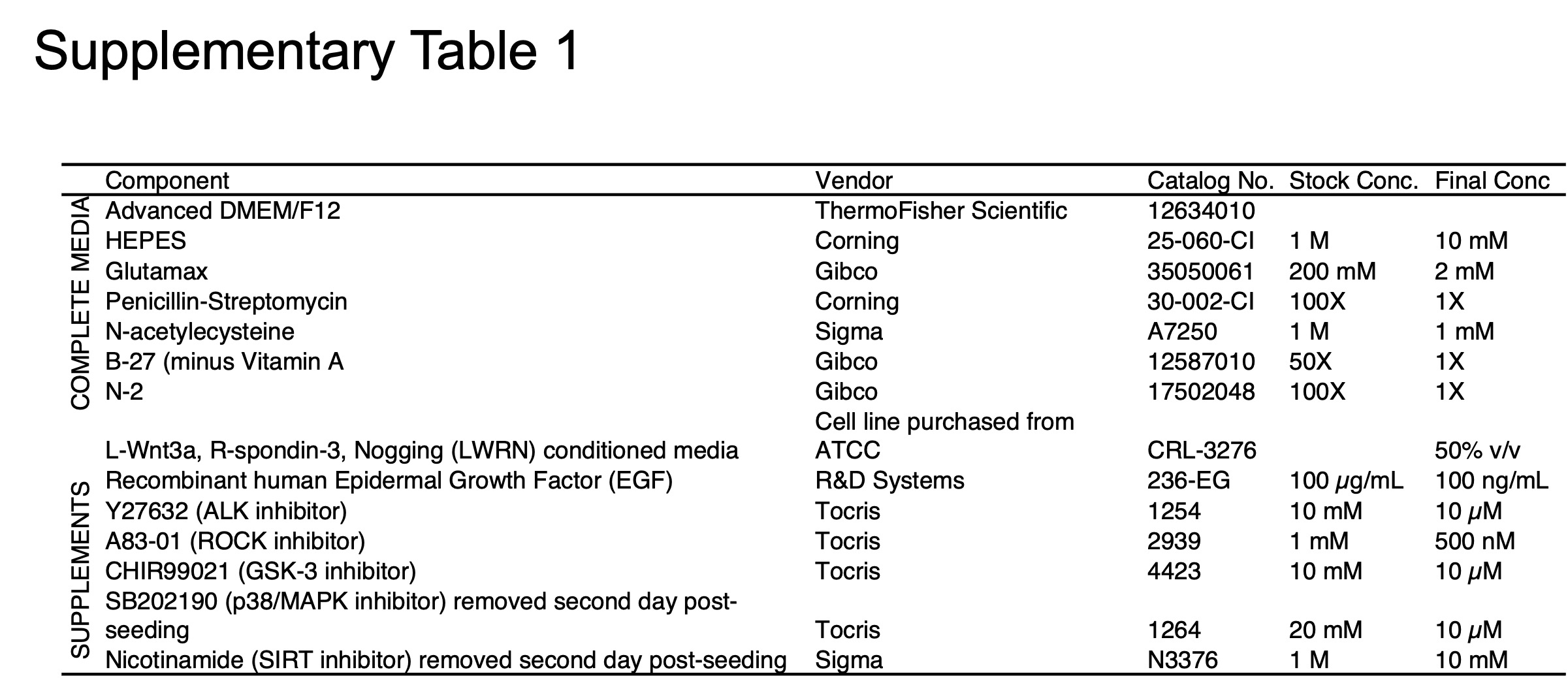
